# Transcriptional dynamics orchestrating the development and integration of neurons born in the adult hippocampus

**DOI:** 10.1101/2023.11.03.565477

**Authors:** Natalí B. Rasetto, Damiana Giacomini, Ariel A. Berardino, Tomás Vega Waichman, Maximiliano S. Beckel, Daniela J. Di Bella, Juliana Brown, M. Georgina Davies-Sala, Chiara Gerhardinger, Dieter Chichung Lie, Paola Arlotta, Ariel Chernomoretz, Alejandro F. Schinder

**Author notes:** equally contributing authors (Natalí B. Rasetto, Damiana Giacomini, Ariel A. Berardino). equally contributing authors (Paola Arlotta, Ariel Chernomoretz, Alejandro F. Schinder).

## Abstract

The adult hippocampus generates new granule cells (aGCs) that exhibit distinct functional capabilities along development, conveying a unique form of plasticity to the preexisting circuits. While early differentiation of adult radial glia-like neural stem cells (RGL) has been studied extensively, the molecular mechanisms guiding the maturation of postmitotic neurons remain unknown. Here, we used a precise birthdating strategy to follow newborn aGCs along differentiation using single-nuclei RNA sequencing (snRNA-seq). Transcriptional profiling revealed a continuous trajectory from RGLs to mature aGCs, with multiple sequential immature stages bearing increasing levels of effector genes supporting growth, excitability and synaptogenesis. Remarkably, four discrete cellular states were defined by the expression of distinct sets of transcription factors (TFs): quiescent neural stem cells, proliferative progenitors, postmitotic immature aGCs, and mature aGCs. The transition from immature to mature aCGs involved a transcriptional switch that shutdown molecular cascades promoting cell growth, such as the SoxC family of TFs, to activate programs controlling neuronal homeostasis. Indeed, aGCs overexpressing *Sox4* or *Sox11* remained stalled at the immature state. Our results unveil precise molecular mechanisms driving adult neural stem cells through the pathway of neuronal differentiation.

## INTRODUCTION

The hippocampus is involved in processing spatial representations and plays multiple functions in regard to memory acquisition, storage and retrieval ^1,2^. The dentate gyrus is the primary gateway for information coming from the entorhinal cortex to the hippocampus. It bears an architecture with unique dynamics due to the presence of radial glia-like neural stem cells (RGLs), which continuously generate new neurons that integrate in the preexisting networks ^3–5^. Adult-born granule cells (aGCs) provide a substrate for the plasticity of perforant-path to GC synapses (dentate input) as well as for mossy fiber to CA3 connections (output) ^6–9^. Such intensive circuit remodeling is crucial for the fine discrimination of similar experiences ^10–12^.

In the mouse brain, developing aGCs display distinct functional characteristics until they become mature after >8 weeks, a time course that is known to last several months in primates ^3,13^. The maturation of functional properties over time involve reduction of membrane resistance, expression of voltage-gated channels, afferent and efferent synaptogenesis, switch of GABA-mediated signaling from excitation to inhibition (**Figure 1A**). At 4 weeks, aGCs display a transient period of enhanced excitability and susceptibility to activity-dependent synaptic plasticity. This period represents a crucial contribution of neurogenesis to circuit remodeling and information processing in the dentate gyrus ^6,8,9,14–22^. Once mature, aGCs are functionally indistinguishable to GCs born during perinatal development ^23,24^. Each step of neuronal differentiation is precisely shaped by the activity and physiological conditions of local dentate networks. Behaviors that increase the dentate activity such as exercise, environmental enrichment or spatial learning exert positive modulatory effects, while conditions that alter the adult neurogenic niche such as aging, inflammation or neurodegeneration are typically detrimental for neurogenesis ^4^. The molecular mechanisms controlling the transitions throughout aGC development and their modulation remain unknown.

**Figure 1.**
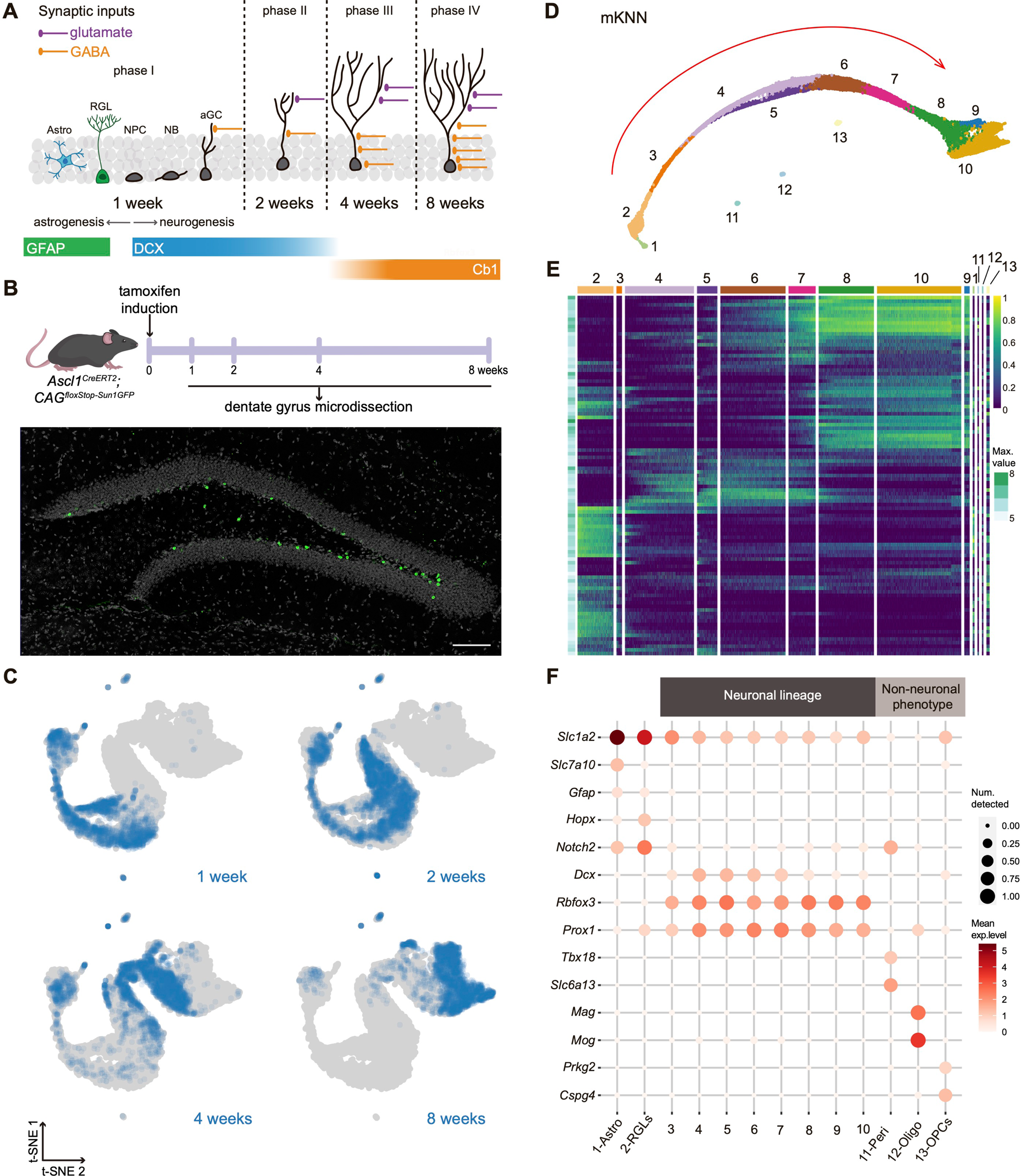
Transcriptional dynamics of neuronal cohorts born in the adult dentate gyrus. **(A)** Schematic representation of developing aGCs in the adult hippocampus. RGLs have the potential to give rise to both astrocytes and neurons. Adult neurogenesis can be divided in four phases. Phase I: radial glia-like neural stem cells (RGLs) generate astrocytes (Astro) or neural progenitor cells (NPCs), which subsequently give rise to neuroblasts and then granule cells (aGCs). New aGCs receive dendritic depolarizing GABAergic inputs. Phase II: developing aGCs acquire glutamatergic contacts. Phase III: period of enhanced excitation/inhibition balance and high excitability. Phase IV: all inputs including perisomatic GABAergic inhibition become mature and excitability decreases ^78^. Colored bars denote the expression of GFAP (astrocytic lineage), DCX and Calb1 (neuronal lineage). **(B)** Experimental design. *Ascl1^CreERT2^*;*CAG ^floxStop-Sun1/sfGFP^* mice received tamoxifen (TAM) injections to label nuclei from the progeny of RGLs and NPCs. Dentate gyri were microdissected at indicated time points. The confocal image below depicts a dentate gyrus with Sun1/sfGFP-labeled nuclei from 2-week-old aGCs. Scale bar: 100 µm. **(C)** T-distributed stochastic neighbor embedding (t-SNE) visualizations for individual neuronal cohorts collected at 1, 2, 4 and 8 weeks after labeling (total: 14,367 nuclei). **(D)** Mutual *k* nearest neighbor (mKNN, *k*=40) graph displaying 13 clusters indicated by color and number. The red arrow denotes the path followed by the process of adult neurogenesis from the least- to the most advanced stages. **(E)** Heatmap showing the normalized expression for 100 genes with highest variability (listed in **Table S1**), which highlights the specific transcriptomic signature for each cluster. Numbers and colors correspond to clusters shown in (D). Maximal expression of each gene is shown on the left (Max. value). Pseudocolor scale on the right denotes mean expression level. **(F)** Dot plot exhibiting canonical markers for neuronal and non-neuronal phenotypes. Pericyte (Peri): *Tbx18* and *Slc6a13*; Oligodendrocyte (Oligo): *Mag* and *Mog*; Oligodendrocyte progenitor (OPC): *Prkg2* and *Cspg4*; astrocyte (Astro): *Gfap, Slc7a10* and *Slc1a2*; RGL: *Hopx* and *Notch2*; neurons: *Dcx, Rbfox3* and *Prox1*. Scales on the right correspond to log2(mean gene expression) (color) and to the fraction of nuclei expressing at least one transcript count in the cluster (dot size). All data in the figure correspond to dataset 1.

Recent studies have exploited the power of single-cell transcriptomics to provide comprehensive descriptions of the initial stages of adult neurogenesis, thus identifying RGLs with distinct potential for self-renewal and differentiation ^25,26^. Additional work has strengthened the long-standing notion that RGLs generate intermediate neural progenitor cells (NPCs) with high proliferating capacity, which then transition to postmitotic neuroblasts ^27^. Later steps of neuronal differentiation have been inferred after studying dentate gyrus cells isolated from postnatal and adult mice ^28^. However, the progression from RGL to mature neuron in the adult hippocampus has never been studied.

Here, we put forward the hypothesis that developmental transitions respond to distinctive molecular programs that are sequentially activated in aGCs. To capture all intermediate stages through the pathway of aGC differentiation, we labeled defined neuronal cohorts at different ages *in vivo* (from 1- to 8-week-old cells) and performed single-nucleus RNA sequencing (snRNA-seq), generating two independent datasets. The high specificity and temporal resolution of these datasets enabled us to reveal the profile of immature and mature aGCs. Unsupervised algorithms identified ten clusters that assembled into a linear developmental trajectory. This continuous pathway was reflected in the transcriptional dynamics of effector genes including pathfinding and cell-adhesion molecules, ion channels, neurotransmitter receptors, and components of the synaptic machinery. Analysis of differential gene expression, pseudotime trajectory, and transcription factors (TFs) revealed critical transitions defining four cellular states: quiescent neural stem cells, proliferative progenitors, postmitotic immature aGCs, and mature aGCs. The passage from immature to mature aGCs involved a transcriptional switch that shut down TFs controlling cell growth and synaptogenesis and turned on the expression of pathways limiting growth and homeostasis. The shutdown of *SoxC* genes emerged as a crucial event to achieve maturation. Supporting this hypothesis, overexpression of *Sox4* or *Sox11* maintained developing aGCs in a persistent immature phenotype. Overall, our work uncovers a molecular continuum underlying neuronal differentiation in the adult hippocampus, assigning specific transcriptomic profiles to the individual stages, revealing master regulator TFs and the effector molecules that convert those programs into neuronal function.

## RESULTS

### Transcriptomic profiling of adult neurogenesis with high temporal resolution

To profile neurogenesis in the adult mouse hippocampus, we utilized a strategy to isolate developing cells derived from RGLs expressing the proneural gene *Achaete-scute homolog 1* (*Ascl1*) ^25,29,30^. Young-adult *Ascl1^CreERT2^*;*CAG^floxStopSun1sfGFP^* mice were used to induce the expression of Sun1-sfGFP in the nuclear membrane of RGLs and permanently label their progeny upon tamoxifen administration (***Figures 1B and S1A***) ^31^. This birthdating tag enabled a comprehensive analysis of the entire process encompassing neuronal differentiation and functional maturation. *A first dataset was built using FACS-sorted nuclei obtained from dentate gyri microdissected at 1-, 2-, 4-, and 8 weeks after induction (cohorts w1 through w8).* High-throughput s*nRNA-seq carried out using Chromium 10X Genomics 3’ end sequencing technology rendered 14,367 profiles with a median of 2,994 genes/nucleus belonging to all four cohorts (**Figure S1B-E**).* Plotting dataset 1 using t-distributed stochastic neighbor embedding (*t*-SNE) revealed that nuclei from the distinct *cohorts* formed a continuous sequence organized by neuronal age (**Figure 1C**). Considering a mutual k-nearest neighbor graph (mKNN, k=40), we used a graph-based unsupervised (Louvain) clustering procedure to group nuclei into communities ^32^. This initial graph-based partition was further refined and revealed 13 clusters (**Figure 1D**). Clusters #1 through #10 seemed to constitute a linear developmental trajectory, while clusters #11, 12 and 13 appeared separated from each other and distant from the main pathway. All clusters displayed individual signatures that supported the definition of the unsupervised partitions (**Figure 1E**). The identity of each cluster was determined by the expression of canonical markers. The majority of cells (about 88% of dataset 1) were distributed in partitions #3 through #10 that belonged to the neuronal lineage and were identified by the expression *Rbfox3* (*NeuN,* panneuronal marker), *Prox1* (aGCs), and doublecortin (*Dcx,* immature neurons; **Figure 1F**). Although astrocytes (cluster #1) and RGLs (#2) are known to be transcriptionally similar, astrocytes were defined by the restricted expression of *Gfap* and *Slc7a10,* together with *Slc1a2, HopX* and *Notch2* that were also expressed in RGLs ^28,33^.

Clusters #11, 12, and 13 encompassed non-neuronal cells that are not described as *Ascl1*^+^ lineage ^29^. These groups included pericytes expressing *Tbx18* and *Slc6a13* (#11), oligodendrocytes expressing *Mag* and *Mog* (#12), and oligodendrocyte precursor cells expressing *Prkg2* and *Cspg4* (#13). Their scarce representation in the entire dataset (<2%) and their separation from the developmental trajectory suggested a transient expression of *Ascl1* in these cell types not related to neurogenesis.

Both the *t*-SNE and mKNN graphs indicated that aGCs follow a developmental continuum encompassing nine stages (**Figure 2A**). While unsupervised clustering relied entirely on genetic profiles, nuclei followed a linear track ordered by neuronal age in the mKNN graphs (**Figures 2B,C and S1F**). Clusters belonging to the neuronal lineage (*Rbfox3^+^, Prox1 ^+^*) followed a sequence containing neural progenitor cells (NPCs), neuroblasts (NB1, NB2), immature neurons (GCimm1, GCimm2), young neurons (GCyoung), and mature GCs (GCmat1, GCmat2). NPCs were recognized by the expression of *Eomes, Top2a, Neil3*, *Cdk6, Lockd, Mcm6,* and *Pola1*, revealing active cell cycle activity (**Figures 2D and S5A**). Interestingly, the presence of *Elavl2, Igfbpl1, Sox4* and *Sox11* indicated that neuronal determination has already occurred in this early state. Neuroblasts lacked genes involved in cell cycle and expressed immature neuronal markers including *Calb2, Dcx, Rgs6, Sox4, Sox11, and Tac2*. Stages that followed in the pathway, GCimm1 and 2, shut down *Calb2 and Elavl2*, and expressed *Dcx, Sox11, Igfbpl1, Rgs6, Camk4, and Chd5*. Interestingly, GCyoung showed diminished expression of immature neuronal markers and enhanced levels of *Calb1, Icam5, Tenm1,* and *Grin2a.* Finally, clusters belonging to mature neuronal phenotypes GCmat1 and 2 had completely shut-off all immature markers and expressed *Ntng1*.

**Figure 2.**
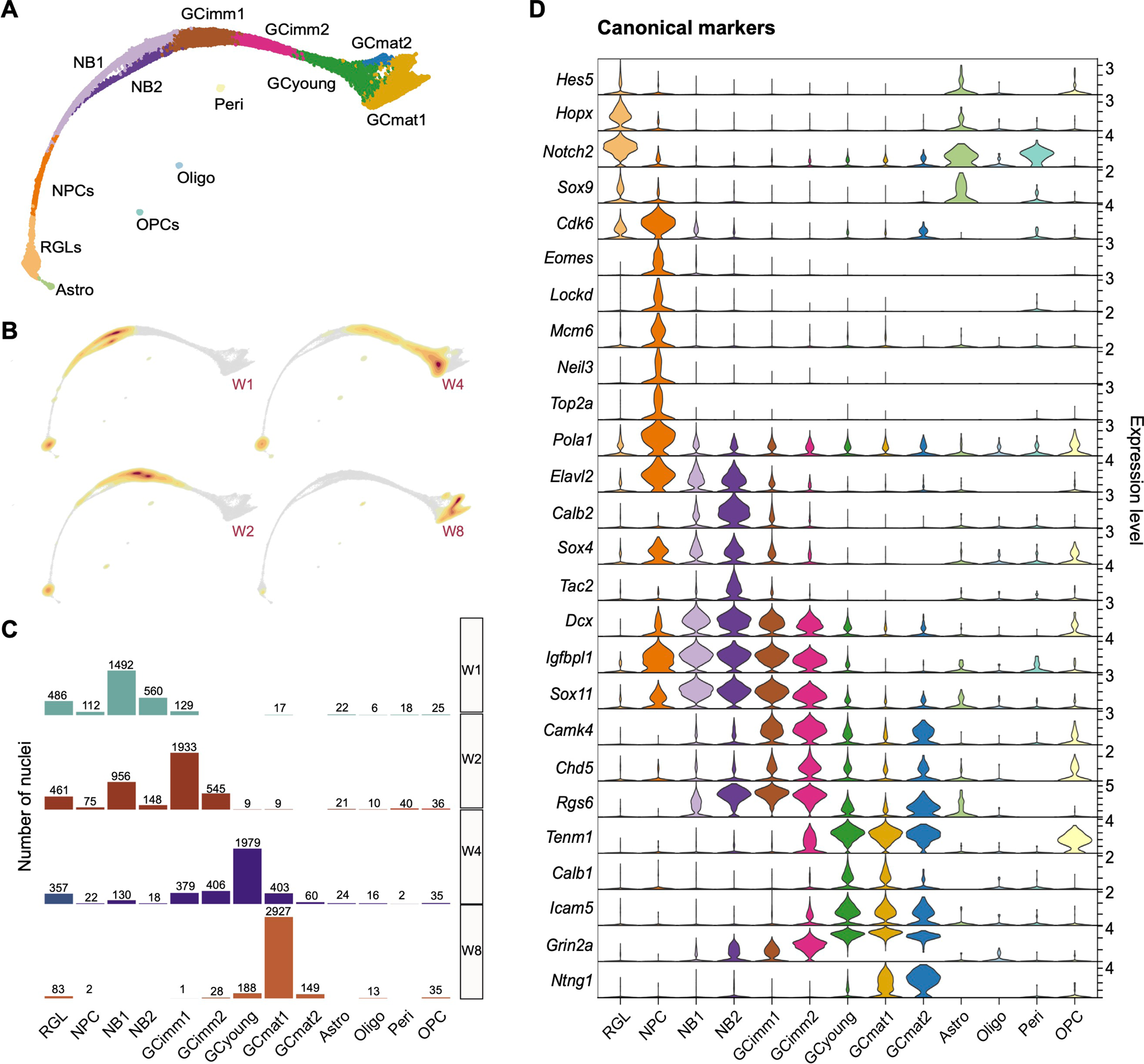
Timing and complexity of adult neurogenesis revealed by molecular profiling. **(A)** mKNN graph displaying cluster identity based on the expression of canonical genes: RGL (radial glia-like cell), NPC (neuronal progenitor cell), NB (neuroblast), GCimm (immature aGC), GCyoung (young aGC), GCmat (mature aGC), Astro (astrocyte), OPC (oligodendrocyte progenitor cell), Oligo (oligodendrocyte) and Peri (pericyte). **(B)** Nuclei distribution for each cohort along clusters. **(C)** Progression of each cohort and their localization over the mKNN graph. Nuclei density is indicated by the yellow (low) to red (high) gradient. **(D)** Violin plots showing the expression level of canonical marker genes for the defined clusters. All data in the figure correspond to dataset 1.

The cohort analysis allowed to define the time course for cluster onset (**Figures 2C and S1F**). Thus, RGLs were detected at early timepoints and declined sharply in w8, in line with the limited proliferative capacity and self-renewal of *Ascl1*-expressing stem cells ^25,30^. NPCs were scarce, as expected based on their fast division and differentiation, and mostly present in w1 and w2. Neuroblasts were distinguished by their early onset and rapid disappearance, particularly NB2. Immature and young aGCs appeared sequentially with distinctive temporal progression. GCimm1 primarily emerged in w1 and reached a maximum in w2, while GCimm2 was observed in w2 and w4. GCyoung was uniquely found in w4, which strongly suggests that this cluster corresponds to developing aGCs undergoing the critical period of enhanced plasticity. GCmat1 was primarily observed in w8. Although dataset 1 was arranged in a continuous pathway with significant changes in cluster composition from w2 to w4 and w8, additional transitions occurring at intermediate times might have been oversighted. To improve time resolution, we collected dataset 2 with cohorts at 2, 3, 4, 5, and 8 weeks (w2 through w8; **Figures S2A,B**). After assessing for the biological reproducibility between both datasets, we used Seurat functionality to transfer cluster labels from dataset 1 to nuclei belonging to dataset 2 (**Figures S3A,B;** see Methods). Clusters in dataset 2 maintained overall properties including differentially expressed genes through transitions, timing of appearance of the distinct clusters, and expression of canonical markers, although with a somewhat slower pace (**Figures S2C,D and S3C**). No nuclei were assigned as NB2, probably because dataset 2 did not include a timepoint at 1 week. GCyoung was again dominant in w4 and GCmat1 was maximally expressed at w8 but it was also present at w5, suggesting an earlier onset.

This transcriptomic analysis describes the continuous trajectory of a unique neuronal type, the adult-born granule cell, through distinct stages from RGL to GCmat. These multiple successive clusters denote higher complexity than the predicted from functional studies, and establishes the molecular basis for the morphological and physiological changes occurring during aGC differentiation.

### Distinct cellular states in the trajectory from neural stem cell to mature neuron

The sequential pathway from RGL to NPC and then NB1 involved profound cell biological transitions, from a quiescent to a proliferating state and then to a postmitotic neuroblast. These transitions displayed the highest numbers of differentially expressed genes (DEGs), particularly in terms of transcript shutdown. In addition, the passage from GCimm2 to GCyoung also exhibited a high number of DEGs with marked gene downregulation (**Figures 3A and S4A**). To better understand the dynamics of the developmental pathway, we assigned pseudotime values to all nuclei along the continuous trajectory from RGL to GCmat1 (**see Methods section**) and organized the datasets to obtain one density plot for each cohort (**Figures 3B and S4B**). Pseudotime density plots displayed four peaks separated by clear valleys. These valleys were coincident with the largest transitions resulting from the DEG analysis.

**Figure 3.**
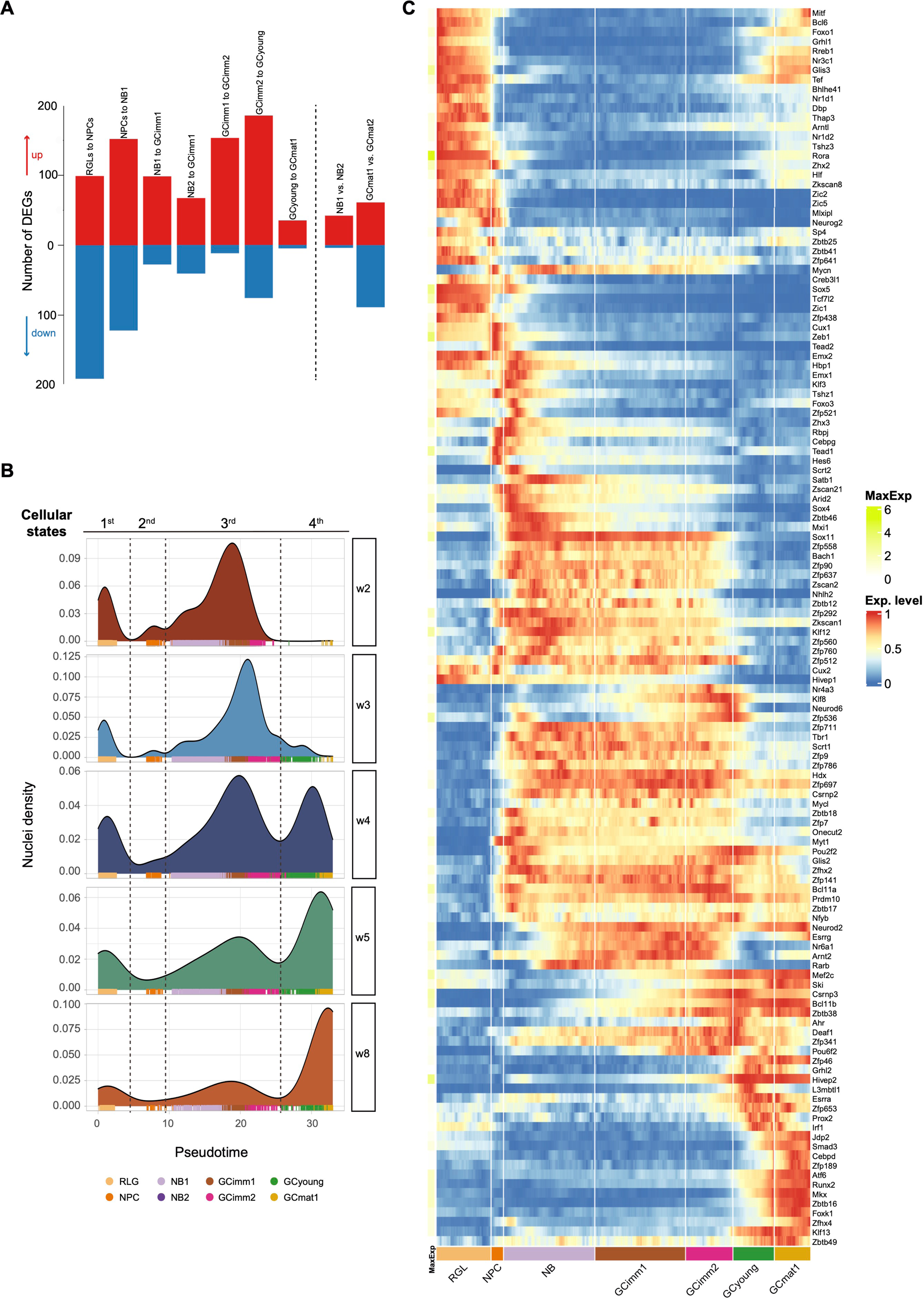
A developmental trajectory defined by four cellular states. **(A)** Differentially expressed genes (DEGs; **Table S2**) between adjacent cluster transitions indicated above (dataset 1). Additional comparisons (non-adjacent) are shown on the right. Up- and downregulated genes are shown in red and blue. **(B)** Density distribution of nuclei along the pseudotime progression for each cohort in dataset 2. Pseudotime values were assigned based on their profile and organized in density plots. The data revealed four cellular states with distinct cluster composition: 1^st^ RGL; 2^nd^ NPC; 3^rd^ NB1-GCimm1-Gcimm2; 4^th^ GCyoung-GCmat1. Dashed lines depict major transitions in gene profile expression. Color codes below denote cluster identity. **(C)** Heatmaps displaying the row-wise normalized expression of transcription factors with sharp transitions switching ON/OFF across clusters. Genes are ordered by the pseudotime of their switching event. Maximum expression (MaxExp) is shown on the left (colored scale on the right). Color-coded clusters are indicated below.

To identify the molecular principles guiding this process, we analyzed the profile of TF expression. Aligning TFs in accordance to their expression onset or shut-off revealed four distinct signatures that were readily observed in the resulting heatmaps: one containing RGLs, another for NPCs, a longer window encompassing all immature neurons (NB1 through GCimm2), and a last interval including GCyoung and GCmat1 (**Figures 3C and S4C**). TF expression displayed different patterns, with a majority restricted to the RGL cluster (such as *Etv4, Etv5, Hes5, Gli1*), others remaining in NPCs (*Ascl1, Sox6, Sox9, Fezf2*), and some covering only the NB stage (*Emx1, Klf3, Zhx3*). Multiple TFs covered the trajectory from NB to GCimm2 (*Sox4*, *Sox11*, *Tbr1*, *Klf12*), while others remained silent through development and became expressed in GCyoung (*Atf6, Runx2, Mkx, Foxk1*). Interestingly, a subset of TFs expressed in RGLs shut down in NPCs and was upregulated again in GCyoung and GCmat1 (*Foxo1, Glis3, Bcl6*, *Hlf*). Within each segment, cell clusters displayed little modifications in TFs. The radical switches in TF expression, the marked changes in the number the DEGs, and the valleys separating those segments in the pseudotime plots revealed three crucial transitions underlying the developmental trajectory. Together, these features defined four distinct cellular states: one corresponding to RGLs, another containing NPCs, a third state enclosing immature neurons from NB1 to GCimm2, and a fourth one encompassing GCyoung and GCmat1.

### Building neuronal function through effector molecules

The transition from RGL to NPC switched off about 200 genes that remained silenced thereafter, reflecting a transformation from a pluripotent program towards a state of active cell division (**Figure 4A,B**). NPCs were the only dividing cluster, expressing multiple cell-cycle genes and displaying an early neuronal commitment revealed by the expression of *Eomes, Sox4, Sox11* and *Igfbpl1* (**Figures 2D and S5A**). The trajectory was followed by two types of neuroblasts, NB1 and NB2, lacking cell-cycle genes but with maximal levels of *Dcx* and *Igfbpl1* expression (**Figure 4C**). The transition from NPC to NB1 upregulated transcripts controlling neuronal differentiation and morphogenesis, cell-cell interaction, development and guidance of neuronal projections, and synaptic organization (**Figure 4D**).

**Figure 4.**
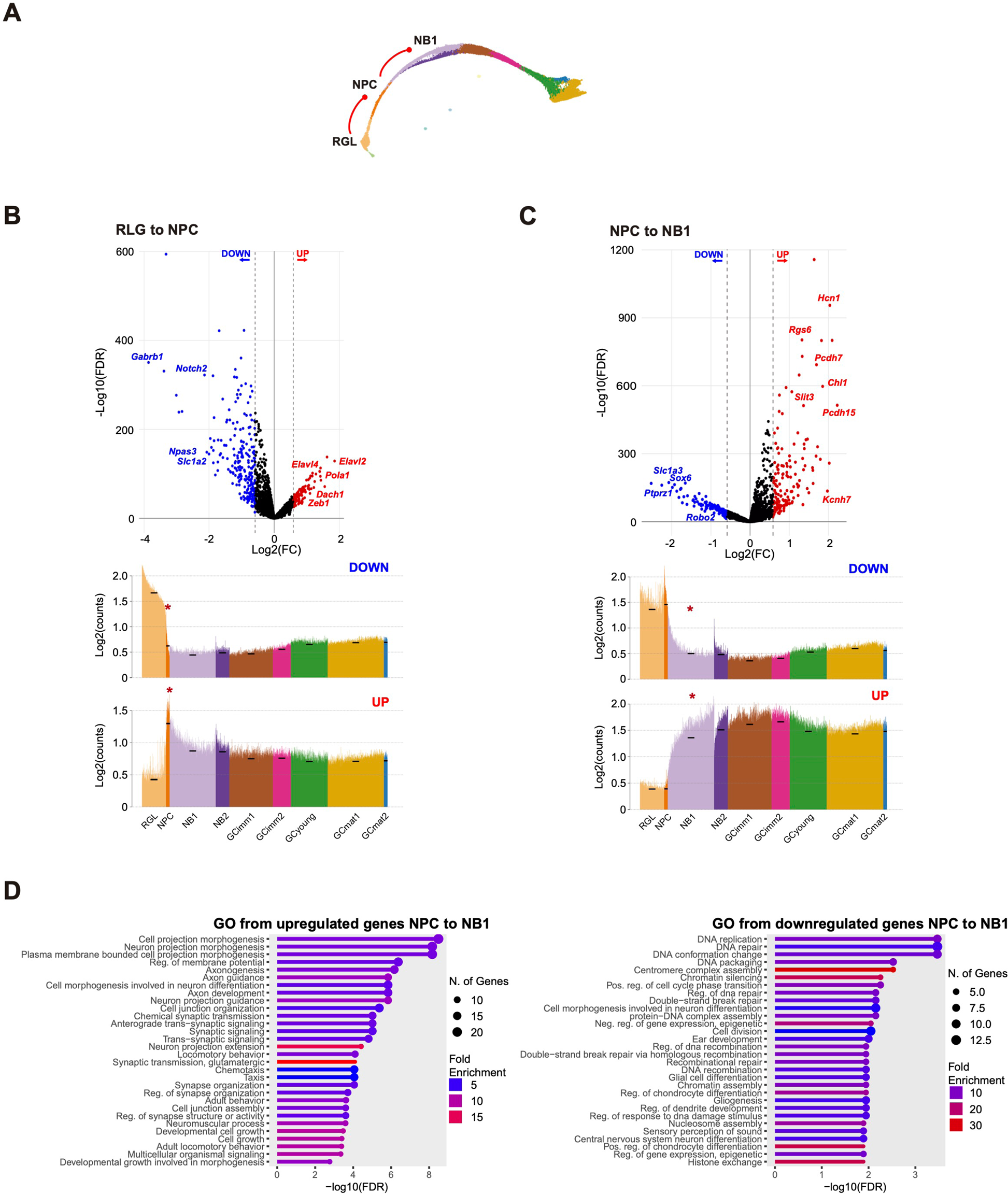
Major switches in gene profiles and biological processes during early neuronal commitment. **(A)** Schematic mKNN graph depicting the analyzed transitions (red lines). **(B, C)** Analysis of RGL to NPC **(B)** and NPC to NB1 transitions **(C).** Upper panels: volcano plots showing differential expression analysis (FC ≥ 1.5 or ≤ −1.5 and FDR ≤ 0.05) displaying relevant gene examples. FC: fold change. FDR: false discovery rate. Lower panels: spike plots displaying mean expression of down- and upregulated DEGs for all nuclei ordered according to their estimated pseudotime value. Black dashes show the mean expression for each cluster and the red asterisks highlight the analyzed transitions. **(D)** Top GO biological processes for the enrichment analysis of DEGs in the transition from NPC to NB1. FDR cutoff = 0.05. All data in the figure correspond to dataset 1. DEGs are listed in **Table S2.**

The canonical markers of early neural development, *Dcx*, *Sox4*, *Sox11*, *Igfbpl1* and *Rgs6*, were expressed in the segment of the trajectory spanning the third state, from NB1 to GCimm2 (**Figures 2D and S3C**). Transitions from neuroblast to GCimm2 primarily involved upregulated gene expression with few downregulated transcripts, suggesting that aGCs continued to incorporate molecules that expanded their structural and functional capabilities (**Figures 3A and S4A**). In general, effector genes that showed an onset of expression in NB1 increased steadily and were maintained until maturation (**Figure 4C**). This was the case for genes related to synaptic transmission such as *Cacna1e, Camk4, Cdh8, Fgf14, Gabrb1, Gabrg3, Grin2a, Grm5, Grm7, Kcnma1* and *Syt1* (**Figure 5A,C,D**). A fraction of the genes upregulated in NB1 remained expressed only in immature neurons, following a pattern similar to the TF signature expressed during the third cellular state (**Figures 3C and S5B,C**). The transition from NB1/NB2 to GCimm1 displayed a marked increase in GABA and glutamate receptor subunits, voltage-gated ion channels and cell-adhesion molecules, among the multiple transcripts supporting neuronal function. Genes related to axon guidance (*Robo1*, *Nrp1*, *Epha7*, *Dscam*, *Ncam2*, *Epha6*, *Slit3*, *Ptpro*) also initiated their expression during the immature stages.

**Figure 5.**
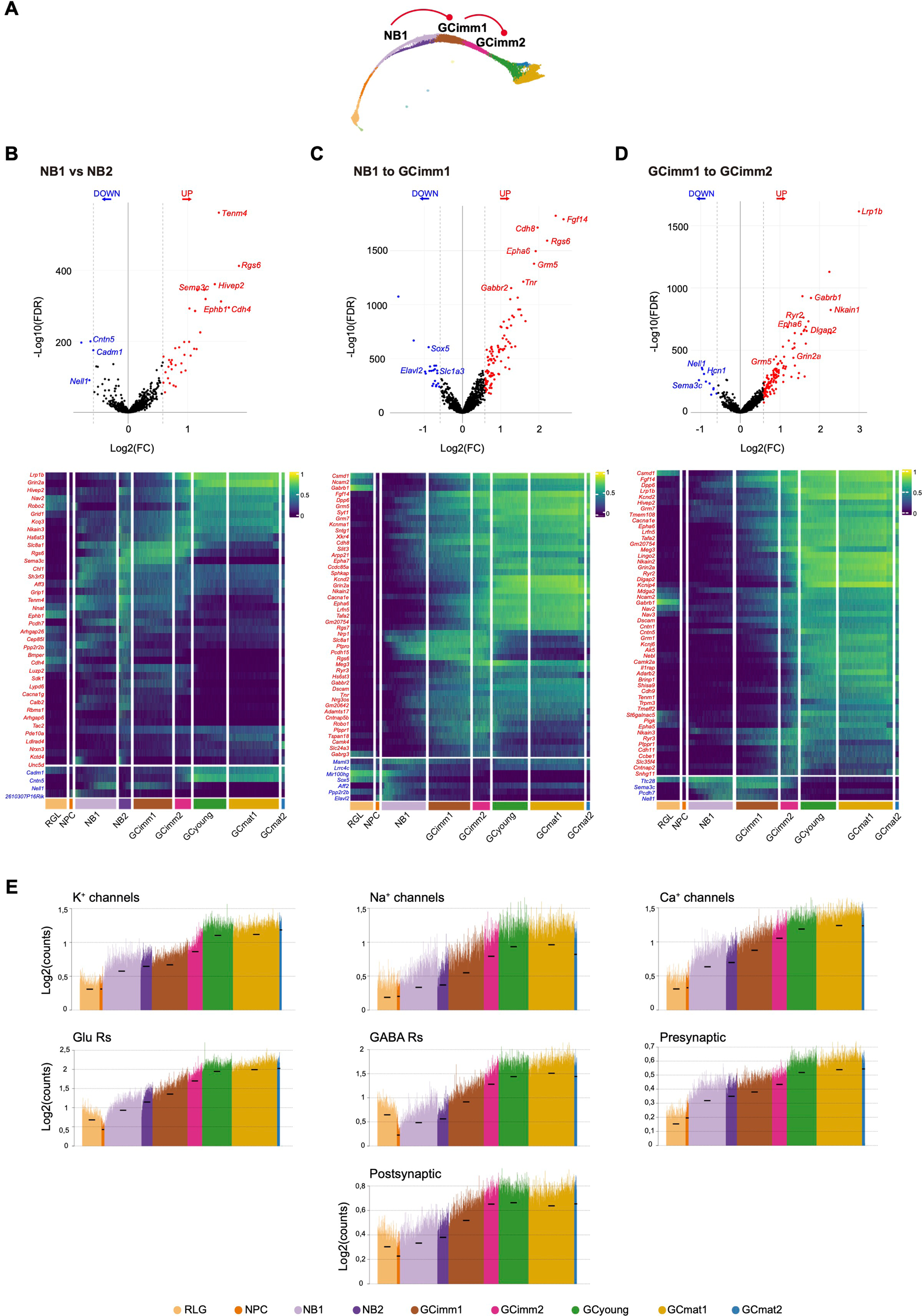
Upregulation of effector genes required for neuronal function in immature aGCs. **(A)** Schematic mKNN graph depicting the analyzed transitions (red lines). **(B, C, D)** NB1 vs NB2 comparison, **(B)** analysis of NB1 to GCimm1 **(C)** and GCimm1 to GCimm2 transitions **(D).** Upper panels: volcano plots showing differential expression analysis (FC ≥ 1.5 or ≤ −1.5 and FDR ≤ 0.05) with relevant gene examples. Lower panels: heatmap showing the row-wise normalized expression of up- (red) and downregulated DEGs (blue) across clusters (top 50 genes per transition). Pseudocolor scale on the right denotes mean expression level. DEGs are listed in **Table S2**. **(E)** Spike plots showing the mean expression of genes relevant for neurotransmission (**Table S4**). Nuclei were ordered according to their estimated pseudotime value. Black dashes show the mean expression for each cluster. All data in the figure correspond to dataset 1.

NB2 constituted a parallel pathway diverging from NB1 towards GCimm1. In the progression from w1 to w2, the NB2 population dropped twice as fast as NB1, suggesting a shorter half-life (**Figures 2C and S1F**). The canonical marker for the neuroblast stage *Calb2* was mainly present in NB2, similarly to *Tac2* and *Rgs6* (**Figure 2D**). Compared to NB1, NB2 displayed about 50 additional transcripts with almost no reduction in gene expression (**Figures 3A and 5B**). Several transcripts continued to increase in GCimm1 and 2, suggesting that NB2 might be at a more advanced developmental stage closer to GCimm1 than NB1. Some of these upregulated genes are related to the assembly of glutamate receptors (*Grin2a, Grid1*, *Grip1*), axon guidance (*Robo2, Sema3c* and *Unc5d*), and cell-cell interaction (*Ephb1, Cdh4, Pcdh7, Nrxn3*)(**Figures 5E and S5D**).

GCimm1 and GCimm2 shared specific markers that were previously described for immature GCs ^28^ (**Figure S6A,B**). Some of these markers, *Adamst18, Nell1, Sema3c* and *Robo1,* were validated by fluorescence *in situ* hybridization (**Figure S6C**). Their expression was confined to the inner granule cell layer, where immature aGCs are typically found ^34,35^. Besides these general similarities, GCimm2 displayed a distinct profile indicating a more advanced developmental stage, including genes related to synapse formation and plasticity, such as *Camk2a* and multiple glutamate receptor subunits: *Grm1*, *Grid1*, *Gria3*, *Grin2a*, *Grm5* and *Grm7* (**Figure 5C,D**)^36–38^. These findings suggest that GCimm2 initiates a program for glutamatergic synaptogenesis which, in electrophysiological studies, was shown to begin in 2-week-old aGCs.

The early clusters described above, RGL, NPC and neuroblasts, reflect different cellular states that bring neural stem cells towards postmitotic neuronal differentiation. In the transitions occurring within the third state (NB to GCimm1 and then to GCimm2), developing aGCs incorporate molecules that expand their functional capabilities, with little transcript shutdown.

### Molecular blueprint of neurotransmission and activity

The passage from GCimm2 to GCyoung revealed a major switch that marked the end of the immature period and the beginning of the fourth state (**Figures 3 and 6A,B**). This transition exhibited a large number of DEGs, highlighting the complexity required to reach neuronal maturation. Multiple transcripts that had reached maximal expression during early development, shut down in GCyoung: *Adamts18, Dcx, Hcn1, Igbpl1, Robo1, Rgs6, Sema3c, Sema3e, Sox4, Sox11*. To better understand the functional changes involved in this transition, we performed enrichment analysis of GO terms (**Figure S7**). Genes related to axonal growth and development shut down, while those orchestrating postsynaptic organization, transmission and plasticity appeared during this period. Furthermore, GCyoung showed plateau levels in typical effector genes related to excitability and neurotransmission (**Figure 5E**). These transcripts included the vesicular glutamate transporter VGLUT1 (*Slc17a7*), the K+/Cl- cotransporter KCC2 responsible for the GABA switch (*Slc12a5*), and *Camk2a* and *Grin2a*, crucial for synaptic plasticity (**Figure 6B**). These features are entirely consistent with prior electrophysiological characterizations of hyperexcitable four-week-old aGCs ^14–16,18,20^.

**Figure 6.**
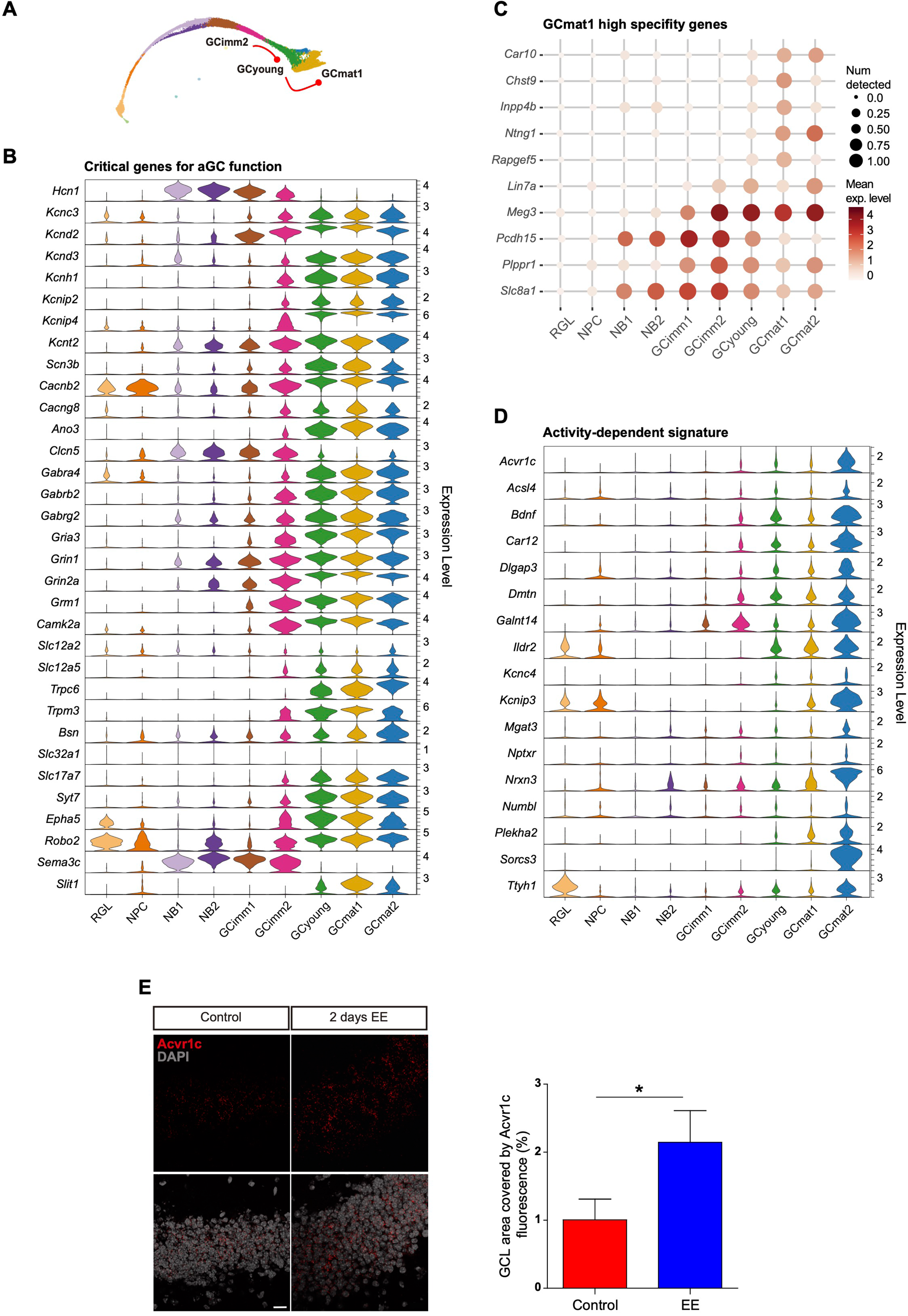
Molecular blueprint of mature aGCs. **(A)** Schematic mKNN graph depicting the analyzed transitions (red lines). **(B)** Violin plots showing example genes critical for aGC function. **(C)** Dot plot exhibiting genes selectively up- and downregulated in GCmat1. Scales on the right correspond to mean expression levels (color) and to the fraction of nuclei expressing at least one transcript count in the cluster (dot size). **(D)** Violin plots depicting the expression of transcripts that are known to be induced by activity (Jaeger et al 2018). All data in the figure correspond to dataset 1. **(E)** *In situ* hybridization reveals the expression of the activity-induced gene *Acvr1c* in the granule cell layer. Mice were exposed to an enriched environment for 2 days (EE) or remained in a home cage (control). Left panels display representative single plane images depicting *Acvr1c* expression (red; DAPI, grey). Scale bar: 20 µm. Right panel: dentate gyrus area occupied by Acvr1c fluorescence in control and EE mice. N = 10 sections / 4 mice (control) and N = 15 sections / 4 mice (EE). (*) denotes *p*<0.02 after Mann Whitney’s test. Data depicts mean±SEM.

The GCyoung and GCmat1 clusters, corresponding to the fourth state, were observed at the w4 and w8 timepoints, respectively, which is coincident with the interval at which aGCs become functionally mature ^3^. This transition was characterized by a few DEGs with subtle changes in their expression level (**Figure S8A,B**). A small group of the DEGs constituted markers of GCmat1: *Car10, Chst9, Inpp4b, Ntng1, and Rapgef5* (**Figure 6C**). Other genes were specifically downregulated in GCmat1: *Lin7a*, *Meg3*, *Pcdh15, Plppr1*, and *Slc8a1.* Notably, a subgroup of cells that expressed genes compatible with a ventral localization, which is associated to different functional properties, was only found in these two clusters (**Figure S8C**)^39,40^. Even though 4- and 8-week-old aGCs are known to be functionally distinct, GCyoung and GCmat1 clusters displayed similar expression profiles. This observation indicates that physiological maturation is also influenced by additional factors other than their transcriptomic signature.

GCmat2 was a small but sharply defined cluster that contained cells belonging to cohorts w4 onwards in both datasets, and exhibited a high number of DEGs when compared to GCmat1 (**Figure 3A**). GCmat2 displayed a set of well-defined markers whose expression has been linked to activity of dentate granule cells: *Acvr1c, Sorcs3, Nrxn3,* and *Kcnip3,* among others ^41^ (**Figure 6D**). Moreover, *Bdnf*, a neurotrophin whose synthesis and secretion depend on neuronal activity, was only expressed in this cluster ^42,43^. Also, *Rgs6* (a marker for immature neurons) has been reported to increase in aGCs after running, and it is upregulated in GCmat2 ^44^. To confirm this activity-dependent signature, mice were exposed to an enriched environment for 2 days, a condition that enhances neuronal activity in the granule cell layer ^45^. This stimulus resulted in a marked increase in the expression of the *Acvr1c* receptor monitored by immunofluorescent *in situ* hybridization (**Figure 6E**). Therefore, GCmat2 was classified as a cluster containing neurons belonging to the w4–w8 timepoints that were likely to be electrically active before isolation.

### A transcriptional switch underlying neuronal maturation

The transitions from RGL to NPC, and NPC to neuroblast were defined by distinct biological programs that enabled switching from quiescent to proliferative and from proliferative to postmitotic states, respectively. The last passage from immature to mature neuron also revealed a major transformation in the transcriptional program. To better understand the biological significance of terminal differentiation, we evaluated the expression of TFs and their target genes (regulons) corresponding to the third and fourth cellular states, aiming to identify master transcriptional regulators for this final step in the developmental trajectory (**Figure 7A**). This analysis (performed using *SCENIC,* see Methods) revealed >20 regulons displaying selective expression in the third state, with highest levels for the *SoxC* family genes *Sox4* and *Sox11.* Regulons present in the fourth state included high levels of *Foxo1*, *Atf6*, *Klf9*, *Tef* and *Hlf* (**Figure S9**). Remarkably, all of these TFs have been shown to control signaling pathways regulating neuronal homeostasis ^46–49^. *Sox4* and *Sox11* have been shown to play a critical role in early differentiation and neuronal growth, both during perinatal development and in adult neurogenesis ^50,51^. However, their participation during late neuronal maturation remains unknown. The shutdown of these *SoxC* TFs in the last neuronal state suggested two possibilities: 1) they are not needed for the final transition and their downregulation results as a secondary effect of a global transcriptional program; 2) their absence might be required for developing neurons to transition to the mature state.

**Figure 7.**
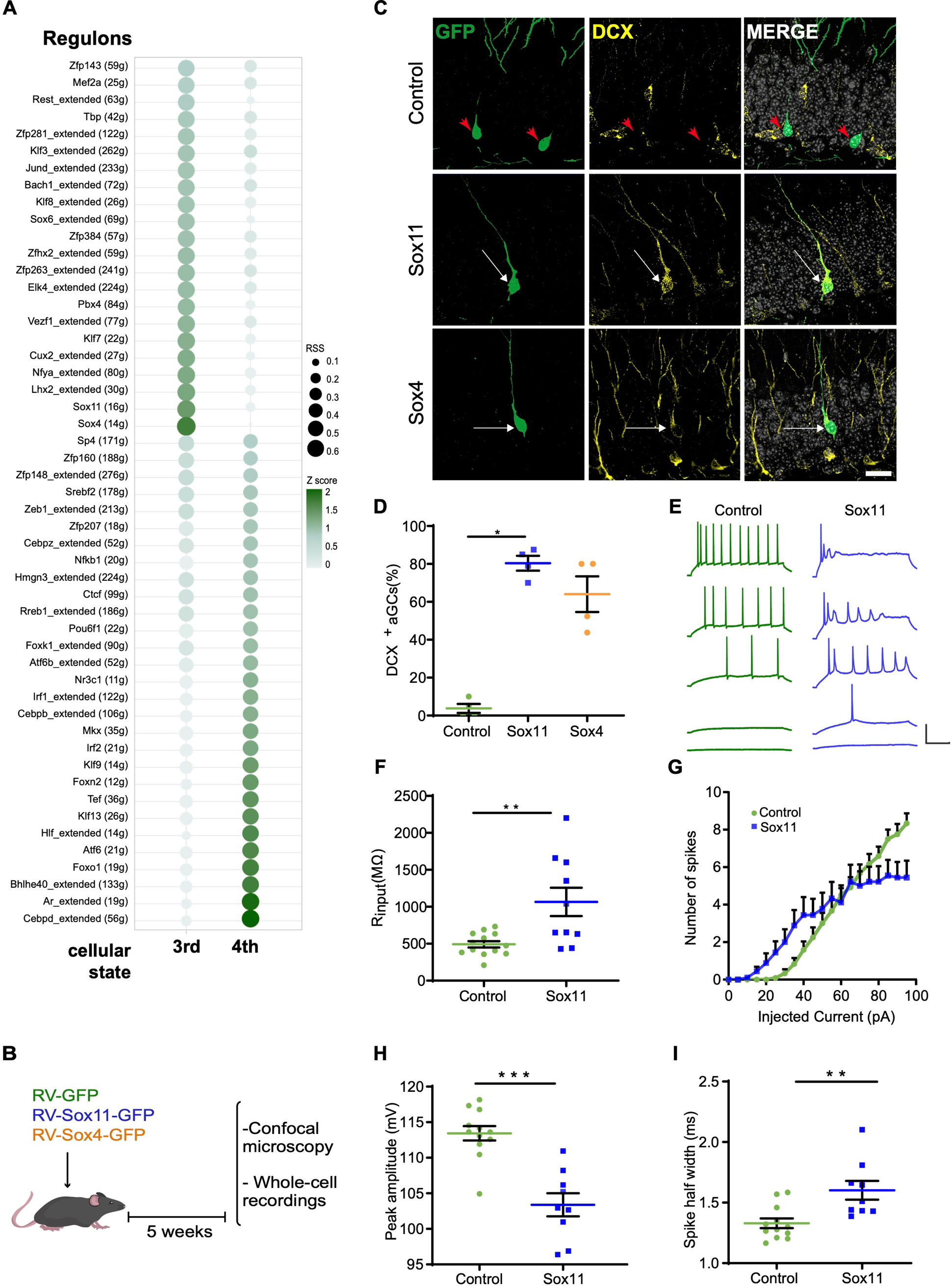
Overexpression of SoxC transcription factors precludes aGC maturation. **(A)** Regulon activity analysis based on the expression levels of TF targets comparing the 3rd and 4th states (dataset 2). Each regulon contains targets genes identified by SCENIC according to the conserved DNA binding motifs in regulatory regions. Regulon sizes are shown in parentheses. Extended regulons include targets inferred by binding motif similarity. Regulon compositions are listed in **Table S5.** RSS: Regulon specificity score ^76^. The z score color scale depicts standardized expression activity values. **(B)** New aGCs were labeled in adult mice by stereotaxic delivery of RV-GFP, RV-Sox11-GFP or RV-Sox4-GFP, as indicated. aGCs expressing GFP were studied 5 weeks post injection. **(C)** Representative single confocal planes displaying aGCs expressing Sox11-GFP, Sox4-GFP or control GFP (green, arrowheads) and DCX (yellow). Colocalization is highlighted by white arrows. Scale bar: 20 μm. **(D)** Proportion of GFP^+^ neurons expressing DCX, normalized to the total number of GFP^+^ neurons. N = 80 aGCs from 4 mice for each condition. (*) denotes *p*<0.02, Kruskal-Wallis test. **(E-I)** Whole-cell recordings in acute slices from aGCs expressing RV-GFP or RV-Sox11-GFP. **(E)** Representative voltage traces elicited by current steps of increasing amplitude: 5, 20, 40, 60 and 95 pA. Scale: 50 mV, 100 ms. V_resting_ kept at −80 mV. **(F)** Input resistance. GFP-aGCs N = 13; Sox11-aGCs, N = 10 (4 mice/condition). (**) denotes *p*<0.005, *t*-test. **(G-I)** Action potentials elicited by injected current steps (500 ms). Plots show the total number of spikes per pulse **(G)**, the amplitude of the first spike **(H)**, and the spike half width, measured at half-maximal amplitude **(I)**. Control GFP-aGCs N = 11 and Sox11-aGCs N = 9 (4 mice/condition). (**) and (***) denote *p*<0.005 and *p*<0.001. Data are shown as mean ± SEM.

To discriminate between these possibilities, we investigated the effect of the sustained expression of *Sox4* or *Sox11* on aGC differentiation. RV-Sox4-GFP or RV-Sox11-GFP were delivered to the dentate gyrus of adult mice to infect mitotic NPCs, coincident with the onset of endogenous expression of *Sox4* and *Sox11* (**Figure 7B**). The impact of TF overexpression in the neuronal progeny was studied after 5 weeks, at which time new aGCs become mature. As revealed by immunofluorescence imaging, the immature neuronal marker DCX was absent in control aGCs but it was observed in 60-80% of neurons overexpressing Sox4 or Sox11, suggesting that cells failed to downregulate DCX upon Sox4/11 overexpression (**Figure 7C,D**). In addition, electrophysiological recordings in acute hippocampal slices showed that aGCs overexpressing *Sox11* displayed high membrane resistance and excitability, as well as impaired capacity to fire repetitive action potentials (**Figure 7E-I**). Together, these results indicate that aGCs with sustained expression of Sox4 or Sox11 maintain immature features ^52,53^. We therefore conclude that shutdown of *SoxC* TFs is a necessary condition for developing aGCs to become mature. The transition from the immature to the mature state constitutes a critical transformation driven by a strong and unique transcriptional switch that shuts down molecular cascades promoting cell growth and activates a final state of neuronal homeostasis.

## DISCUSSION

The molecular basis underlying the differentiation and functional integration of aGCs have remained largely unknown. Previous work has captured aspects of adult neurogenesis by means of transcriptomic profiling with a focus on the early molecular cascades that distinguish quiescent from active RGLs and NPCs ^26,54^. Subsequent research has conveyed the notion of a linear trajectory reaching the early neuroblast stages ^27^. More recently, a study reported that immature GCs obtained from the perinatal, postnatal and adult hippocampus display similar transcriptional profiles ^28^. In this work we performed a comprehensive study encompassing differentiation and functional maturation of aGCs. The strategy involved the analysis of cohorts of aGCs at six timepoints that allowed us to assign precise time stamps to the developmental trajectory. In general, the dynamics of gene expression profiles matched very closely with previous descriptions of developing aGCs obtained using immunohistochemical, morphological and electrophysiological approaches. These datasets now provide a thorough description of distinctive molecular states underlying neuronal differentiation in the adult hippocampus, assigning specific profiles to the intermediate stages, revealing TFs controlling the process, and the effector genes that convert those programs into neuronal function.

The different phases of adult neurogenesis were previously characterized by the expression of stage-specific markers defined by immunolabeling. RGLs gave rise to NPCs and, subsequently, to neuroblasts that were thought to keep limited proliferative capacity ^55,56^. Instead, our results reveal that NPCs are the only mitotic cluster in aGC differentiation, and neuroblasts are committed postmitotic cells ^28^. Two types of neuroblasts were observed: NB1 arising from NPCs, and NB2 shedding from NB1, forming a parallel pathway reaching GCimm1. Several transcripts were upregulated in NB2 and continued to be expressed until GCimm2, which suggests that NB2 might be a more advanced stage of maturation. GCimm1 and GCimm2 were composed mostly of 2- to 4-week-old aGCs and expressed immature neuronal markers. The transition to GCimm2 upregulated >100 genes, including those related to glutamatergic transmission, which is known to begin by the second developmental week ^34,57–59^. This finding indicates that GCimm2 may be the molecular transition that accompanies the onset of glutamatergic synaptogenesis.

Approaching the GCyoung stage involved substantial profile changes (250-350 DEGs in both datasets) that were only comparable in magnitude to the transitions from RGL to NPC, and NPC to NB1. Multiple effector genes reached plateau levels at GCyoung, including those required for neurotransmission and plasticity, from the presynaptic vesicular glutamate transporter *Slc17a7* and vesicular release *Syt7* to postsynaptic glutamate and GABA receptors. Because GCyoung contained predominantly w4 cells, we propose that aGCs in this cluster correspond to 4-week-old aGCs with enhanced excitability and synaptic plasticity, described extensively in the literature ^14–16,18,20,58^. Despite containing the fundamental building blocks required for neuronal function, 4-week-old GCs are not yet mature. In fact, mature aGCs (>8 weeks) exhibit maximal glutamatergic synaptic strength, mature GABAergic inhibition, and reduced excitability ^18,52,60,61^. Few transcripts were found to be selectively changed in the transition from GCyoung to GCmat1, suggesting that the GCyoung cluster is the early phase in a final step of maturation that becomes consolidated over time.

Analysis of DEGs, pseudotime trajectory, and TF expression revealed three critical transitions defining four cellular states: RGL, NPC, immature neurons (NB1 to GCimm2), and a mature state (GCyoung and GCmat; **Figure 8**). Regulon analysis revealed that the immature state was dominated by *Sox4* and *Sox11* regulons, posing them as critical organizers of the regulatory network. These TFs were known to be involved in the early steps of differentiation during perinatal and adult neurogenesis^50,51^. We now demonstrate that shutdown of *SoxC* TFs is a critical mechanism required for achieving terminal differentiation.

**Figure 8.**
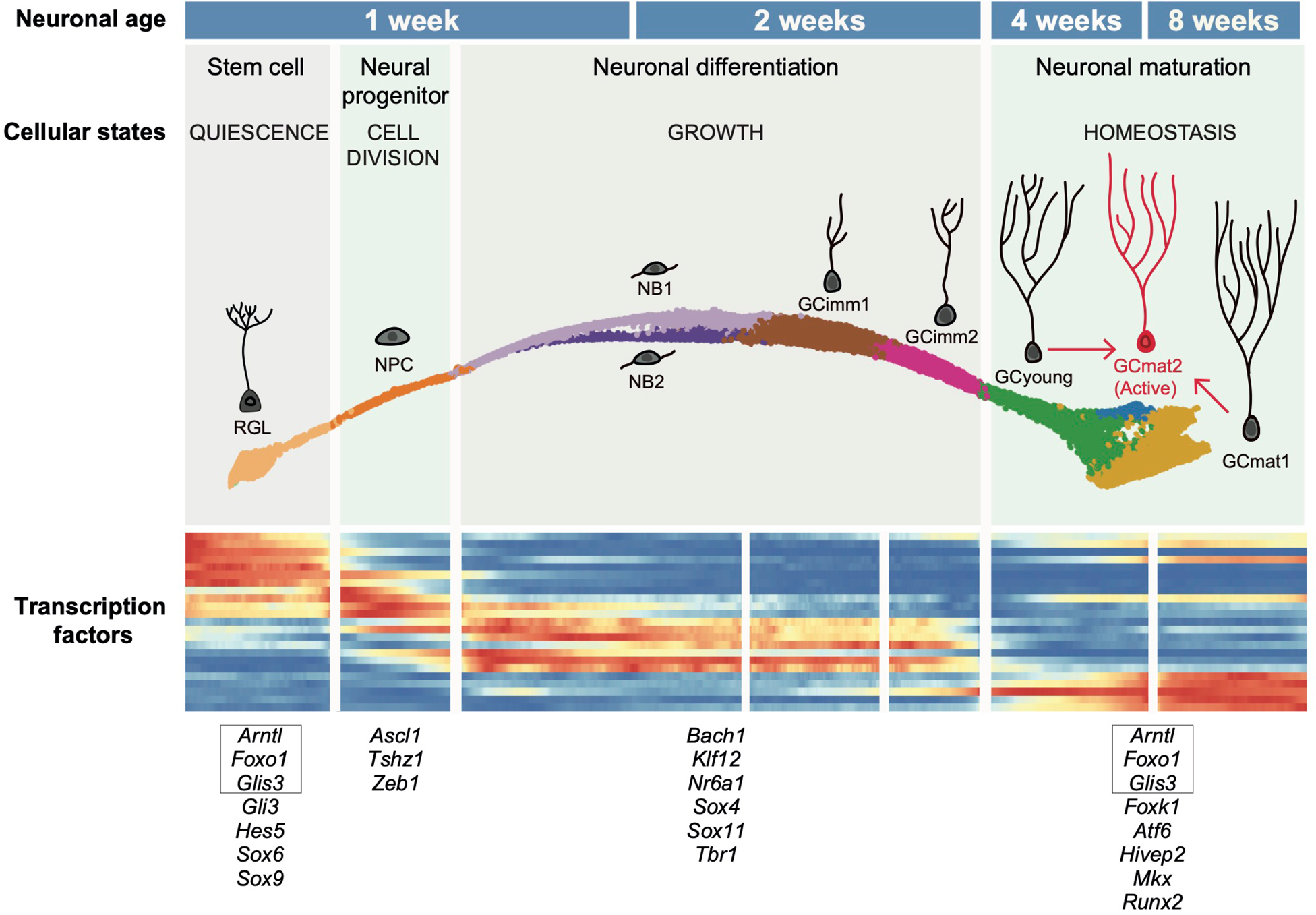
Schematic representation of cell states during aGC maturation. Transcriptomic profiling of developing aGCs unveils a molecular continuum divided in nine clusters grouped into four cellular states, precisely defined by the activation and deactivation of specific transcription factors (TFs). During the developmental period lasting about eight weeks, distinct TFs and effector genes orchestrate the biological processes characteristic of these states. Stem cells abandon the *quiescence* state to become mitotic (*cell division* state). Dividing NPCs rapidly acquire a neuronal fate, entering a state of cellular *growth*. These passages occur during the first developmental week. Postmitotic neurons express a large set of TFs driving differentiation. By the fourth week, growth achieves a plateau and previously expressed programs shut down to activate a set of TFs driving homeostatic control of neuronal function (*homesotasis* state). During this final interval, aGCs are known to become fully integrated in the preexisting network. The heatmap (lower panel) depicts the expression of the example TFs listed below (pseudocolor scale is the same as in Fig. 3C). Squared TFs were expressed both in the quiescent and homeostatic states.

Regulons dominating the fourth state included *Foxo1, Klf9, Hlf, Tef,* and *Atf6*. The known functions of these TFs suggest that they might coordinate a fine homeostatic regulation. For instance, deletion of *Foxo* family TFs has been shown to impair autophagy, leading to aberrant dendritic growth and increased spine density in aGCs ^47^. *Klf9* knockout also resulted in increased spine density and impaired functional maturation in aGCs ^48^. Moreover, genetic ablation of *Hlf* and *Tef* in cultured neurons triggered a disproportionate upregulation of input excitation when deprived of activity ^49^. This evidence points to a central role of these TFs in the homeostatic regulation of excitatory neuronal connectivity. Interestingly, *Atf6* has been associated with calcium homeostasis in models of Huntingtońs disease and endoplasmic reticulum stress ^46,62^. *Foxo1*, *Tef*, *Atf6* and *Mkx* regulons share a common downstream TF, *Hivep2* (Schnurri-2), whose knockout provoked global alterations in the morphology of dentate gyrus GCs ^63^. Finally, *Glis3*, a TF that was selectively expressed in RGLs and then upregulated in GCmat1 (**Figure 3C**), has been associated with autophagy and with the regulation of neuronal growth and complexity ^64^. Together, the regulatory roles of these TFs and their late expression support a hypothesis whereby neuronal maturation is orchestrated by shutting down molecular cascades promoting differentiation, and awaking signaling pathways controlling cellular homeostasis. The data presented here opens a new scenario to unravel the temporal dynamics involving master transcriptional regulators that control neuronal maturation in the healthy adult brain, and will serve as the basis to identify mechanisms contributing to aberrant circuit remodeling in brain disorders.

## ACKNOWLEDGMENTS

We thank members of the A.F.S., A.C., and Emilio Kropff labs for insightful discussions, and Mariela Trinchero for critical comments on the manuscript. D.G., A.C. and A.F.S. are investigators in the Consejo Nacional de Investigaciones Cientificas y Tecnicas (CONICET). N.B.R, A.B. and M.B. were supported by CONICET fellowships. This work was supported by grants from the National Institute of Neurological Disorders and Stroke (NINDS, R01NS128117) to P.A., NINDS and Fogarty International Center (FIC) (R01NS103758) to A.F.S. and P.A., and the Argentine Agency for the Promotion of Science and Technology (PICT 2017-0389) to D.G., (PICT 2018-03713) A.C., and (PICT-2020-0046 and PICT-2021-0077) A.F.S.

## METHODS

### Animals

Male and female C57Bl/6J wild type and genetically modified mice (6–8 weeks of age) were housed at four to five mice per cage in standard conditions. For adult neurogenesis snRNA-seq experiments, *Ascl1^CreERT2^* (*Ascl1 ^tm1(Cre/ERT2)Jejo/J)^*) mice were crossed to *CAG^floxStop-Sun1/sfGFP^* (B6.129*-Gt(ROSA)26Sort^m5.1(CAG– Sun1/sfGFP)Nat/MmbeJ^*) conditional reporter line to generate Ascl1^CreERT2^;CAG^floxStop-Sun1/sfGFP^ mice, which were used to reliably target adult-born GC nuclei ^30,31,65^. Tamoxifen (TAM) induction (120 µg/g, four injections in two consecutive days) resulted in the expression of Sun1-sfGFP in the nuclear envelope of Ascl1^+^ cell progeny. Mice were anesthetized (150 μg ketamine/15 μg xylazine in 10 μl saline per g) and sacrificed at the indicated times after TAM induction: 1-, 2-, 4- and 8 weeks for dataset 1, and 2-, 3-, 4-, 5- and 8 weeks for dataset 2. Mice were maintained in C57BL/6J background. Experimental protocols were approved by the Institutional Animal Care and Use Committee of the Leloir Institute, according to the Principles for Biomedical Research involving animals of the Council for International Organizations for Medical Sciences and provisions stated in the Guide for the Care and Use of Laboratory Animals

### Tissue dissection

Mice were deeply anesthetized (ketamine/xylazine, as described above), brains were carefully removed and placed into ice cold Earl’s balanced salt solution (EBSS, in mM: 117 NaCl, 5.4 KCl, 1 NaH_2_PO_4_, 26 NaHCO_3_, 5.6 glucose, 1.8 CaCl. 2H O and 0.8 MgSO) with trehalose 5% v/v and kynurenic acid 0.8 mM ^66^. Dissection solution was equilibrated in 95% O_2_/5% CO_2_ before use. The brain was cut along the longitudinal fissure and the regions posterior to lambda were cut off. Under a dissection microscope and upon removal of the diencephalon, the medial side of the hippocampus was exposed. The dentate gyrus was isolated by inserting a sharp-needle tip and sliding it superficially along the septotemporal axis of the hippocampus. The dissected tissue was placed in a 0.5 ml microcentrifuge tube with a minimal amount of media, flash-freezed on dry ice and stored at −80°C until use.

### Nuclei isolation and FACS sorting

Nuclei were isolated as previously described with several modifications ^67^. Dounce tissue grinder and pestles were sequentially washed with 100% EtOH, RNAse Zap (Sigma, Cat # R2020), 2-3 rounds with RNAse-free water, and finally rinsed with EZ Lysis Buffer (Sigma, Cat # NUC-101). All material and buffers were chilled on ice. The tissue was transferred to a chilled dounce prefilled with 2 ml ice-cold EZ Lysis Buffer and homogenized slowly with 20-25 strokes of pestle A, followed by 20 strokes with pestle B. Suspension was transferred to a chilled tube and incubated on ice 5 min. Then the suspension was centrifuged at 500 × g for 5 min at 4°C and the supernatant was removed. The pellet was resuspended in 4 ml ice-cold EZ Lysis Buffer and after a second round of ice incubation and centrifugation, the isolated nuclei were resuspended in 4 ml ice-cold Nuclei Suspension Buffer (NSB) (1x RNase-free molecular-biology grade PBS, 0.1% molecular-biology grade BSA 100 μg/ml and 0.2U/μl RNase inhibitor). Finally, nuclei were centrifuged at 500 × g for 5 min at 4°C and after removing the supernatant were resuspended in 1ml NSB-Ruby (final Ruby concentration 1:500) and filtered twice through a 35-μm cell strainer to remove as many debris as possible. Nuclei were kept on ice until sorting (Harvard University, Bauer Core Facility, BD FACSAria II) into 96 well plates, pre-coated with BSA to reduce adherence and improve recovery, containing with ∼10-20 μl of rich-NSB (with final 2U/μl of RNase inhibitor and 1% BSA). The final nuclei concentration was determined using a C-chip Neubauer counting chamber and Trypan Blue (1:2 final dilution). Nuclei were immediately loaded for single-cell GEM formation (10x Genomics, single cell RNA sequencing 31, Chromium v3.1).

### 10x Genomics Chromium

Dataset sampling for snRNA-seq experiments was carried out within a minimal time interval, processing two time points in a single experiment each day to minimize batch effects. For each time point, four dentate gyri (left hemispheres) from male and female mice were pooled. If needed to increase nuclei number, dentate gyri from the right hemisphere were also processed (dataset 2). For snRNA-seq, nuclei were loaded in the 10x Genomics chips aiming to recover 2,000–10,000 nuclei. cDNA amplification and library construction were done following 10x Genomics protocols. For both dataset 1 and dataset 2, libraries were generated using Chromium v3.1, quantified in BioAnalyzer and sequenced on an Illumina HiSeq or NovaSeq. Samples were sequenced to a depth of 40–70,000 reads per cell.

### RNA *in situ* hybridization

For validation of cluster markers, mice were anesthetized and the brains were removed and placed into the ice-cold dissection solution (described above). Hippocampi were dissected under a microscope and immediately embedded in OCT on dry ice and stored at –80 °C. Sections of 14 μm covering the anterior– posterior axis of the dentate gyrus were collected in a cryostat. Sections were placed for 1 h at −20 °C and stored at –80 °C until use. Fluorescent multiplex RNA in situ hybridization was performed using the RNAscope Fluorescent Multiplex Reagent Kit (Advanced Cell Diagnostics), according to the manufacturer’s instructions. Briefly, thawed sections were fixed in PFA 4%, dehydrated in sequential incubations with ethanol, followed by 30 min protease IV treatment. Mice were exposed to the enriched environment for 48h (**Figure 6E**) prior to tissue sample preparation as described above. Appropriate hybridization probes (Advanced Cell Diagnostics, catalogue#: Adamts18 452251-C1, Acvr1c 429291-C1, Nell1 807961-C2, Robo1 475951-C1, Sema3c 441441-C1) were incubated for 2 h at 40 °C, followed by amplification steps (according to protocol), DAPI counterstaining, and tissue was mounted with gerbatol to prevent bleaching.

### Production of viral vectors

A replication-deficient retroviral vector based on the Moloney murine leukemia virus was used to specifically transduce aGCs as done previously ^18,45^. Retroviral particles were assembled using three separate plasmids containing the capside (CMV-vsvg), viral proteins (CMV-gag/pol) and the transgenes: CAG-GFP, CAG-EGFP-IRES-Sox11 and CAG-EGFP-IRES-Sox4. Plasmids were transfected onto HEK 293T cells using deacylated polyethylenimine. HEK 293T cells were cultured in Dulbeccós Modified Eaglés Medium with high glucose, supplemented with 10 % fetal calf serum and 2 mM glutamine. Virus-containing supernatant was harvested 48 h after transfection and concentrated by two rounds of ultracentrifugation. Virus titer was typically ∼10^5^ particles/μl.

### Stereotaxic surgery for retroviral delivery

Running wheel housing was only available 2-3 days before surgery to maximize the number of retrovirally transduced neurons. The wheel was removed one day after surgery. For the stereotaxic procedure, mice were anesthetized and retroviral particles were infused into the dorsal region of the right dentate gyrus (1-1.5 μl at 0.15 µl/min) using sterile calibrated microcapillary pipettes through stereotaxic references. Coordinates from bregma (mm): −2 anteroposterior, −1.5 lateral, −1.9 ventral. A single injection site was sufficient to label an abundant number of neural precursor cells. Brain sections were prepared 5 weeks post infection for immunofluorescence or electrophysiological recordings.

### Immunofluorescence

Immunostaining was performed in 60 μm-thick free-floating coronal sections throughout the septal fraction of the hippocampus from *Ascl1^CreERT2^*;*CAG^floxStop-Sun1/sfGFP^* or retrovirally injected mice. Antibodies were applied in TBS with 3% donkey serum and 0.25% Triton X-100. Immunofluorescence was performed using the following primary antibodies: DCX (rabbit polyclonal; 1:1500; Abcam), GFP (chicken IgY Fraction; 1:500; Aves Labs Inc.). The following corresponding secondary antibodies were used: donkey anti-chicken Cy2 or Cy3 and donkey anti-rabbit Cy5 1:250 (Jackson ImmunoResearch Laboratories). Incubation with DAPI (10 minutes) for nuclear counterstain was performed. Slices were mounted and covered with gerbatol to prevent bleaching.

### Confocal Microscopy

Sections from the septal hippocampus according to the mouse brain atlas (antero-posterior, −0.94 to −2.46 mm from bregma) were included ^68^. Images were acquired using a 880 LSM Airyscan microscope (Carl Zeiss, Jena, Germany). Analysis of antibody expression was restricted to cells with fluorescence intensity levels that enabled clear identification of their soma. For Sox/GFP experiments, images were acquired (40x; NA 1.2) from 60-µm-thick sections, and colocalization was assessed using single optical planes (airy unit=1). For RNAscope experiments, images were acquired from 14 um-thick hippocampal slices. Dot area quantification was performed in ImageJ/Fiji applying the Otsu threshold as suggested by the manufacturer (Advanced cell diagnostics). All experiments were done with control vs. treatment conditions blind to the operator.

### Electrophysiology

Five weeks post retroviral injection, mice were anesthetized and brains were removed into a chilled solution containing (mM): 110 choline-Cl^−^, 2.5 KCl, 2.0 NaH PO, 25 NaHCO, 0.5 CaCl, 7 MgCl, 20 dextrose, 1.3 Na^+^-ascorbate, 0.6 Na^+^-pyruvate, and 4 kynurenic acid. Slices (400-μm thick) were cut in a vibratome (Leica VT1200S) and transferred into a chamber containing artificial cerebrospinal fluid (ACSF; in mM): 125 NaCl, 2.5 KCl, 2.3 NaH PO, 25 NaHCO, 2 CaCl, 1.3 MgCl, 1.3 Na^+^-ascorbate, 3.1 Na^+^-pyruvate, and 10 dextrose (315 mOsm). Slices were maintained with 95% O_2_/5% CO_2_ at 30°C for >1 h before experiments started. Whole-cell recordings were performed at 23 ± 2°C using microelectrodes (3– 5MΩ) pulled from borosilicate glass (KG-33; King Precision Glass) and filled with (mM): 120 K-gluconate, 20 KCl, 5 NaCl, 4 MgCl_2_, 0.1 EGTA, 10 HEPES, 4 Tris-ATP, 0.3 Tris-GTP, 10 phosphocreatine, Alexa Fluor 488 or 594 (10 μg/ml; Invitrogen), pH 7.3, and 290 mOsm. Recordings were obtained using an Axopatch 200B amplifier (Molecular Devices), digitized (Digidata 1322A), and acquired at 10 kHz into a personal computer using the pClamp 9 software (Molecular Devices). Recorded neurons were visually identified in the granule cell layer by fluorescence (FITC fluorescence optics; DMLFS, Leica) and infrared DIC videomicroscopy. Input resistance was obtained from current traces evoked by a hyperpolarizing step of 10 mV. In current-clamp recordings the resting membrane potential was kept at −70 mV by passing a holding current. Criteria to include cells in the analysis was visual confirmation of GFP in the pipette tip, and absolute leak current <50 pA at −60 mV. Series resistance was typically <25 MΩ, and experiments were discarded if >45 MΩ. In all experiments, control or Sox11 overexpression treatments were blind to the operator.

### snRNAseq processing

We used STARsolo 2.7 to align the snRNA-seq reads to the GRCm38 mouse genome. Multi-mapped reads were excluded and both exonic and intronic counts were considered. We used default parameters to count UMI and filter high-quality cells in order to generate gene-by-cell count matrices. Quality control (QC) and downstream data processing were performed using *scran* and *scDblFinder* bioconductor libraries ^69,70^. All scripts used to perform the present analysis were included in the companion website https://github.com/chernolabs/NeuronalSwitch.

For dataset 1, we removed 80 nuclei with <1000 detected features, and 636 nuclei identified as doublets by scDblFinder out of the 17046 nuclei that passed the STARsolo quality filter. We also eliminated 839 low-quality nuclei identified by either a multivariate outlier detection procedure (*adjOutlyingness* function of robustbase R library) based on low library sizes, low number of detected features and large mitochondrial (Mt) content, or 2) presenting more than 1% of Mt counts. Features with >20 molecules and detected in >1% and <80% of filtered nuclei were retained. In addition, we removed specific genes (*Ehd2, Espl1, Jarid1d, Pnpla4, Rps4y1, Xist, Tsix, Eif2s3y, Ddx3y, Uty, Kdm5d, Rpl26, Gstp1, Rpl35a, Erh, Slc25a5, Pgk1, Eno1, Tubb2a, Emc4, Scg5*) to mitigate possible sex and stress effects on downstream analysis ^28^. The data was normalized using the scran library in accordance with the OSCA pipeline ^71^. We executed a quick clustering for each timepoint independently, normalized nuclei in each cluster separately, and rescaled the size factors to be comparable across clusters. Finally, we computed the log2 of the data and considered a pseudo-count value of 1 to generate a normalized expression matrix. After scaling and log-normalizing gene expression values, we used the scran *modelGeneVar* function to identify the 3000 most variable genes in their log-expression profiles.

For dataset 2, 32,621 nuclei successfully passed the STARsolo quality filter. Subsequently, we identified and discarded 1,015 doublets and 4,808 low-quality nuclei using the QC protocol applied to dataset 1. The entire curation process yielded a high-quality dataset comprising 26,798 nuclei and 14,220 informative features. We considered inter-nuclei similarities using the Pearson correlation measure within the subspace spanned by the top 20 principal components. We finally retained for further analysis 26,716 nuclei that were part of graph components larger than 30 nodes. We then used the *batchelor* Bioconductor package to investigate the biological reproducibility between datasets 1 and 2. Some dataset 1 nuclei occupied a region in reduced dimensionality spaces (PCA, t-SNE, UMAP) not represented in dataset 2. Employing a k=20 mKNN-graph estimated from the largest 20 PCA components (similarities obtained from Pearson correlation values), we verified that these nuclei formed two Louvain clusters composed of 1071 nuclei from 4-week-old samples. Noticeably, we found that 85% of nuclei in these clusters exhibited higher expression of male-associated genes (*Ddx3y*, *Uty*, *Eif2s3y*) than female-associated genes (*Xist*, *Tsix*), suggesting that they could belong to a single male mouse, potentially a biological outlier. Therefore, we re-processed *de-novo* dataset 1, excluding these nuclei which resulted in 14,441 high-quality nuclei and 13,353 features. To get a proxy of the biologically relevant manifold, we constructed a mKNN graph (k=40) considering the Pearson correlation similarity measure estimated from the 20 largest PCA components. As we did with dataset 2, we discarded nuclei that were found in small components (<30 elements) of this mKNN graph. The remaining 14,367 nuclei were grouped into communities considering the Louvain algorithm. This initial graph-based clustering was further refined to produce 13 communities for which specific markers were identified (see Results section). We used Seurat’s label transfer functionality to impute cluster labels for data set 2. Only 7 nuclei were identified as NB2, probably due to the fact that dataset 2 did not include a timepoint at 1 week. These nuclei were disregarded for subsequent analysis. We determined differentially expressed genes between nuclei groups using the *FindMarkers* function from the *scran* library (*t*-test pairwise). Cluster marker genes were identified as highly ranked differentially expressed genes between the query group and all other clusters.

### Pseudotime determination

The *slingshot* 2.4.0 R-package was used to fit developmental trajectories to nuclei of the following clusters: RGL, NPC, NB1, NB2, GCimm1, GCimm2, GCyoung and GCmat1 ^74^. We excluded from this analysis GCmat2 nuclei and 383 GCmat1 nuclei with strong expression signal of ventral genes (**FigS8C**). We considered the UMAP (3D) low dimensional coordinates to fit principal curves and produce pseudotime estimations (PCA-3dim and PCA-20dim produced similar density profiles along the corresponding pseudotime coordinates). A similar procedure was used for dataset 2. For the sake of the pseudotime analysis, a cluster refinement of the Seurat label transferred partition was considered, in order to correct visible misclassifications observed in UMAP3D space. This issue affected <3% of nuclei (mainly involving Astrocytes, Pericytes, RGL and NPC nuclei, see 05_DS2_SeuratLT.R in the web companion site).

### TF expression cascades

We considered 700 mouse TFs listed in the *dorothea* 1.8.0 R package that passed the quality control feature filtering step ^72^. We tested whether these genes were significantly expressed along the pseudotime coordinate using the *difftest* function of the TSCAN 1.34 R package. Likelihood ratio tests were performed to compare a generalized additive model with a constant fit to get the p-values, with 201 TFs showing FDR adjusted q-value <0.05. A custom script (09_cascades_Tfs.R in the web companion site) was used to identify TFs that were shutdown (112) or turned on (69) at specific clusters in this dataset (**Figure 3C and S4C**).

### Gene Ontology analysis

GO enrichment analyses was performed using ShinyGO 0.77 (http://bioinformatics.sdstate.edu/go/) ^73^.

### SCENIC regulons

To analyze the transcriptional changes occurring when neurons become mature, we considered 3-, 4-, 5-, and 8-week cohorts of dataset 2 and selected nuclei located around the 3^rd^ and 4^th^ peak of the pseudotime density distribution (19.4±2.5 and 30.3±2.5 pseudotime intervals in **Figure 3B**), containing 4855 and 4244 nuclei respectively. We used the *SCENIC* 1.3.1 R-package to analyze the expression of relevant TFs and their targets (regulons) ^75^. We used the calcRSS function to estimate Regulon Specificity Scores ^76^.

### Visualization and graphical reports

We used Force Atlas 2 layout algorithm, implemented in Gephi to produce 2-dim visualization of mKNN graphs ^77^. Heatmaps, volcano plots, violin plots, dot plots, and spike plots were produced using the scX R package [https://chernolabs.github.io/scX/].

---

**Figure S1.**
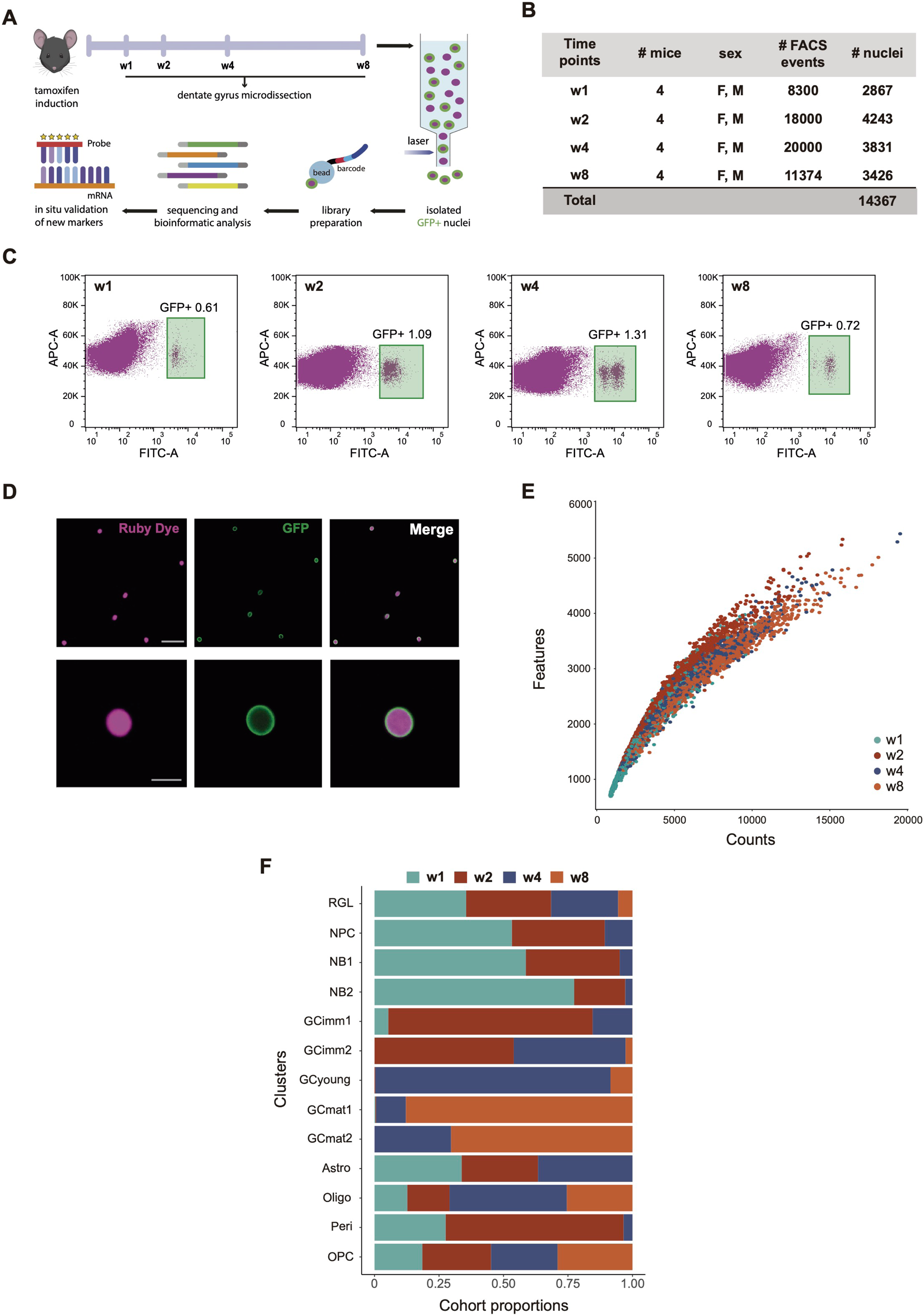
Experimental pipeline and quality control for dataset 1. **(A)** *Ascl1^CreERT2^*;*CAG^floxStop-Sun1/sfGFP^* mice received tamoxifen (TAM) injections to label nuclei from the progeny of RGLs and NPCs. Dentate giri were microdissected at the indicated timepoints (cohorts). Each cohort of GFP^+^ nuclei was FACS-purified and processed separately using 10x Chromium technology for cDNA library preparation and sequencing. Markers were studied by fluorescence *in situ* hybridization. **(B)** Nuclei composition for each cohort, indicating mice number and sex (F, females; M, males), number of FACS-sorted, and analyzed nuclei. **(C)** FACS sorting of Ruby^+^/GFP^+^ nuclei. Scatter plots of ruby dye intensity vs. log(GFP intensity) for purified nuclei isolated at w1, w2, w4 and w8. Green squares highlight the GFP^+^/Ruby^+^ population, with the number indicating % GFP^+^ nuclei in the total population. **(D)** Confocal images of FACS-purified Ruby^+^ nuclei exhibiting GFP anchored to the nuclear membrane. Scale bars: 50 μm (upper panels) and 10 μm (lower panels). **(E)** Quality measurements of snRNA-seq libraries. Scatter plot depicting the number of genes (features) vs. the number of mRNA molecules (counts) per nuclei for each timepoint. snRNA-seq detected similar number of genes and transcript molecules across cohorts. **(F)** Contribution of timed cohorts to cluster composition.

**Figure S2.**
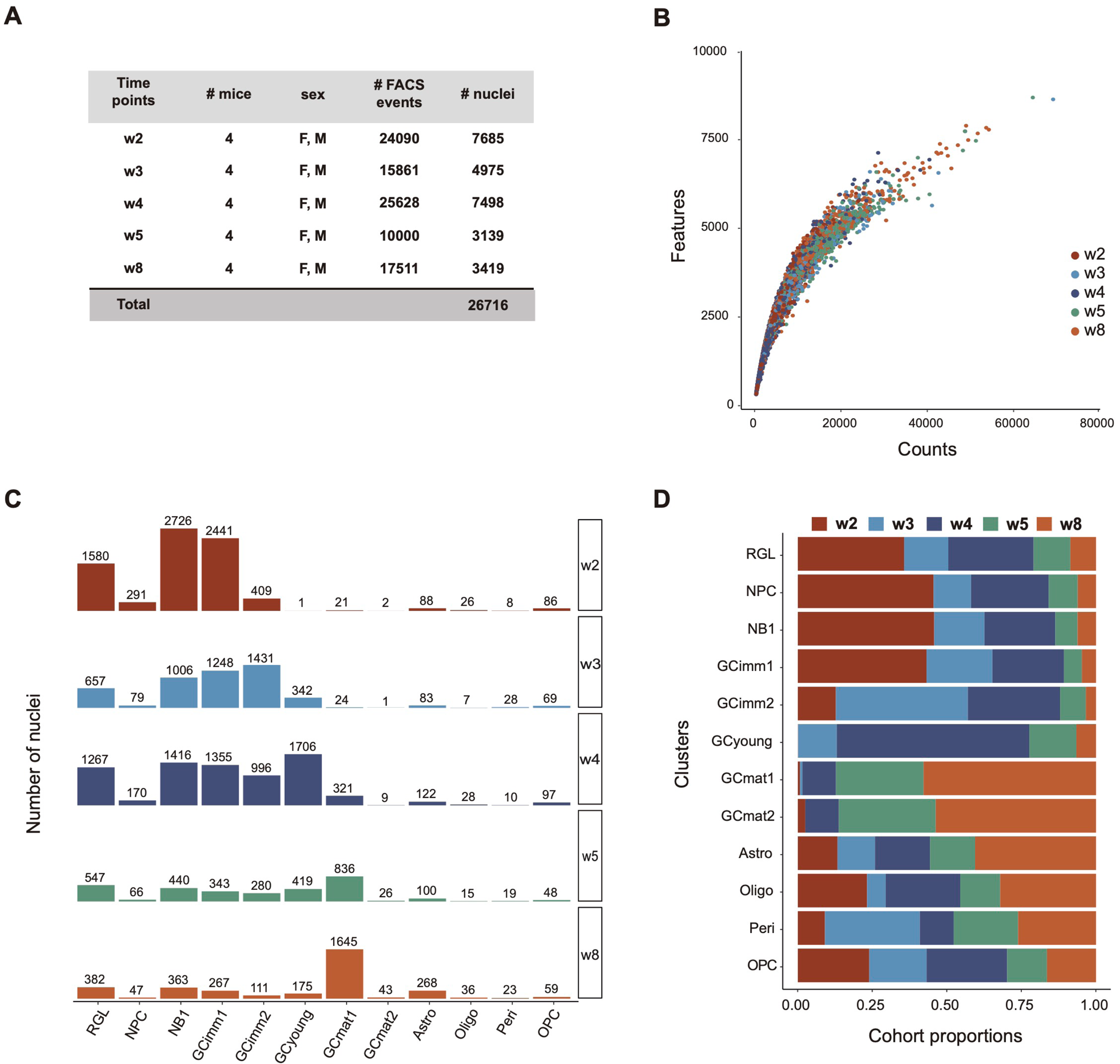
Quality control and cluster distribution for dataset 2. **(A)** Nuclei composition for each cohort, indicating mice number and sex (F, females; M, males), number of FACS-sorted, and analyzed nuclei. **(B)** Quality measurements of snRNA-seq libraries. Scatter plot depicting the number of genes (Features) vs. the number of mRNA molecules (counts) per nuclei for each timepoint. snRNA-seq detected similar number of genes and transcript molecules across cohorts. **(C)** Nuclei distribution in all clusters for each neuronal cohort. **(D)** Contribution of timed cohorts to cluster composition.

**Figure S3.**
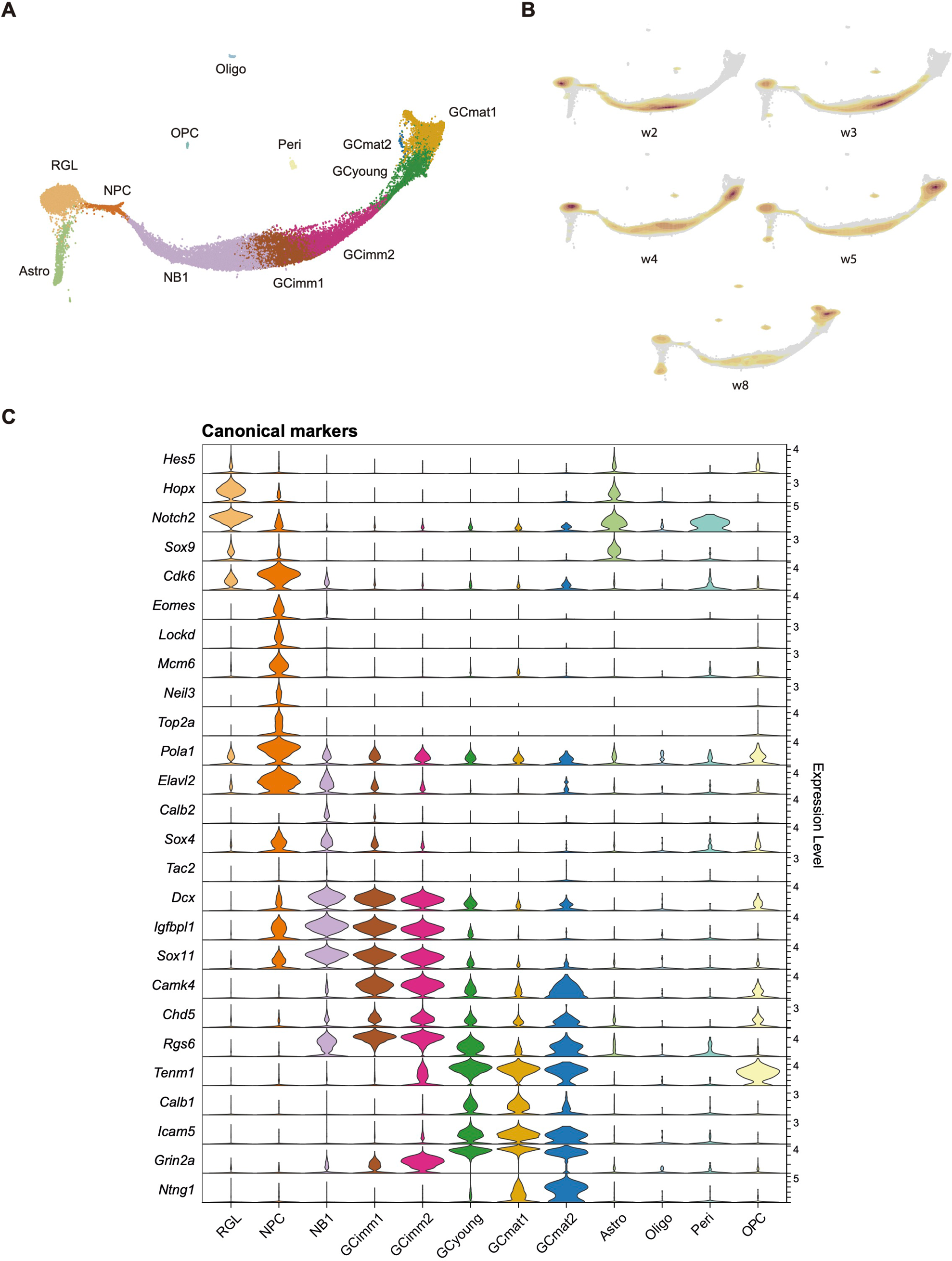
Cluster identity and transitions in dataset 2. **(A)** mKNN graph displaying cluster identity. **(B)** Progression of each cohort and their localization over the mKNN graph. Nuclei density is indicated by the yellow (low) to red (high) gradient. **(C)** Violin plot showing the expression level of canonical marker genes for the defined clusters. Note that most identified clusters are conserved with dataset 1, highlighting the reproducibility between experiments.

**Figure S4.**
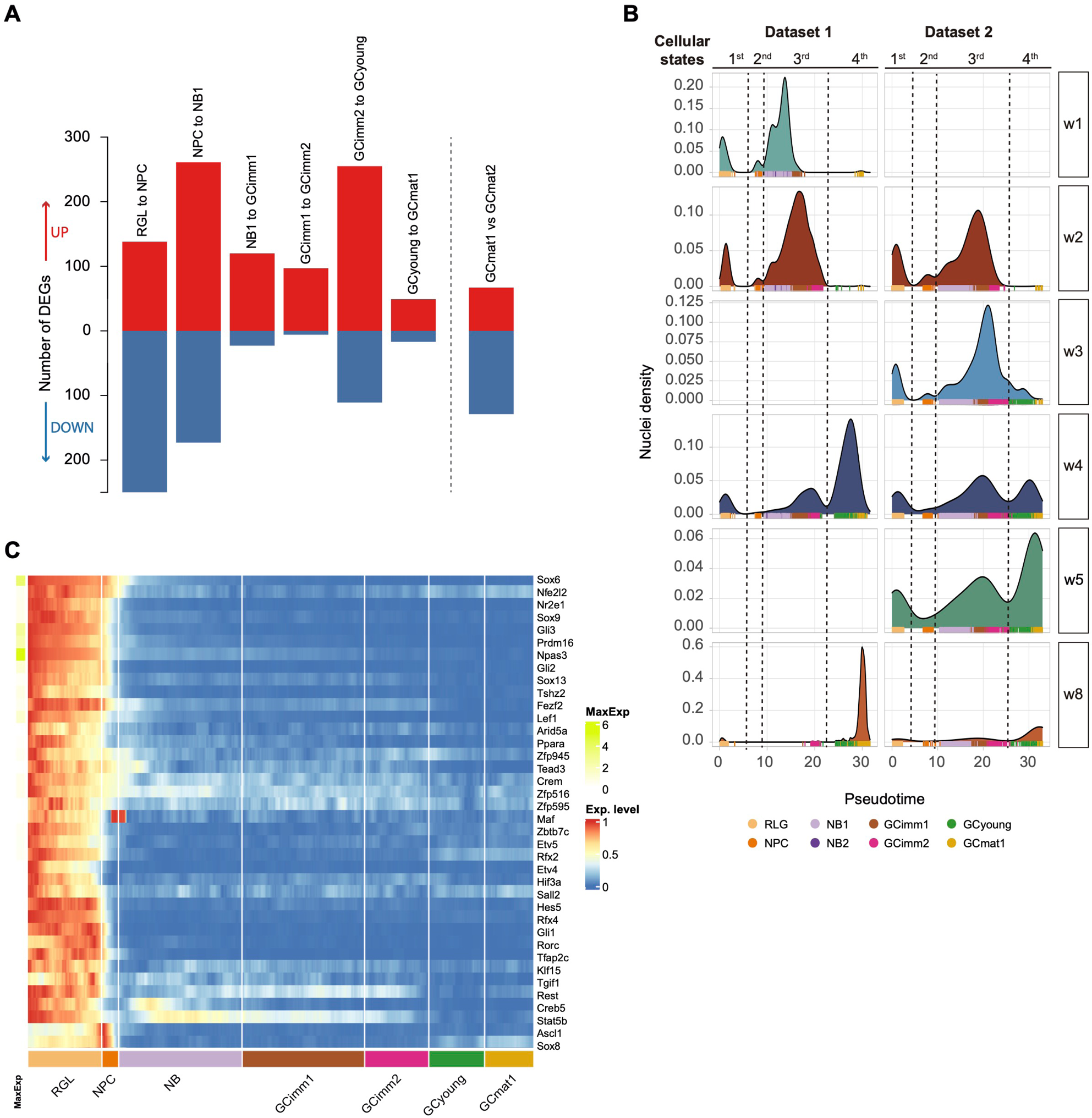
Pseudotime and density profiles are conserved between datasets. **(A)** DEGs between adjacent cluster transitions indicated on top (dataset 2; DEGs listed in **Table S3**). An additional comparison (non-adjacent) is shown on the right. Up- and downregulated genes are shown in red and blue. **(B)** Density distribution of all nuclei along the pseudotime progression for each cohort, comparing datasets 1 and 2. The four cellular states are conserved: 1^st^ RGL; 2^nd^ NPC; 3^rd^ NB1-GCimm1-GCcimm2; 4^th^ GCyoung-GCmat1. Dashed lines depict major transitions along development. Color codes below denote cluster identity **(C)** Heatmap displaying the row-wise normalized expression of transcription factors specific for RGLs, depicting additional transcripts to those shown in Fig. 3C. Maximum expression (MaxExp) is shown on the left (colored scale on the right). Color-coded clusters are indicated below.

**Figure S5.**
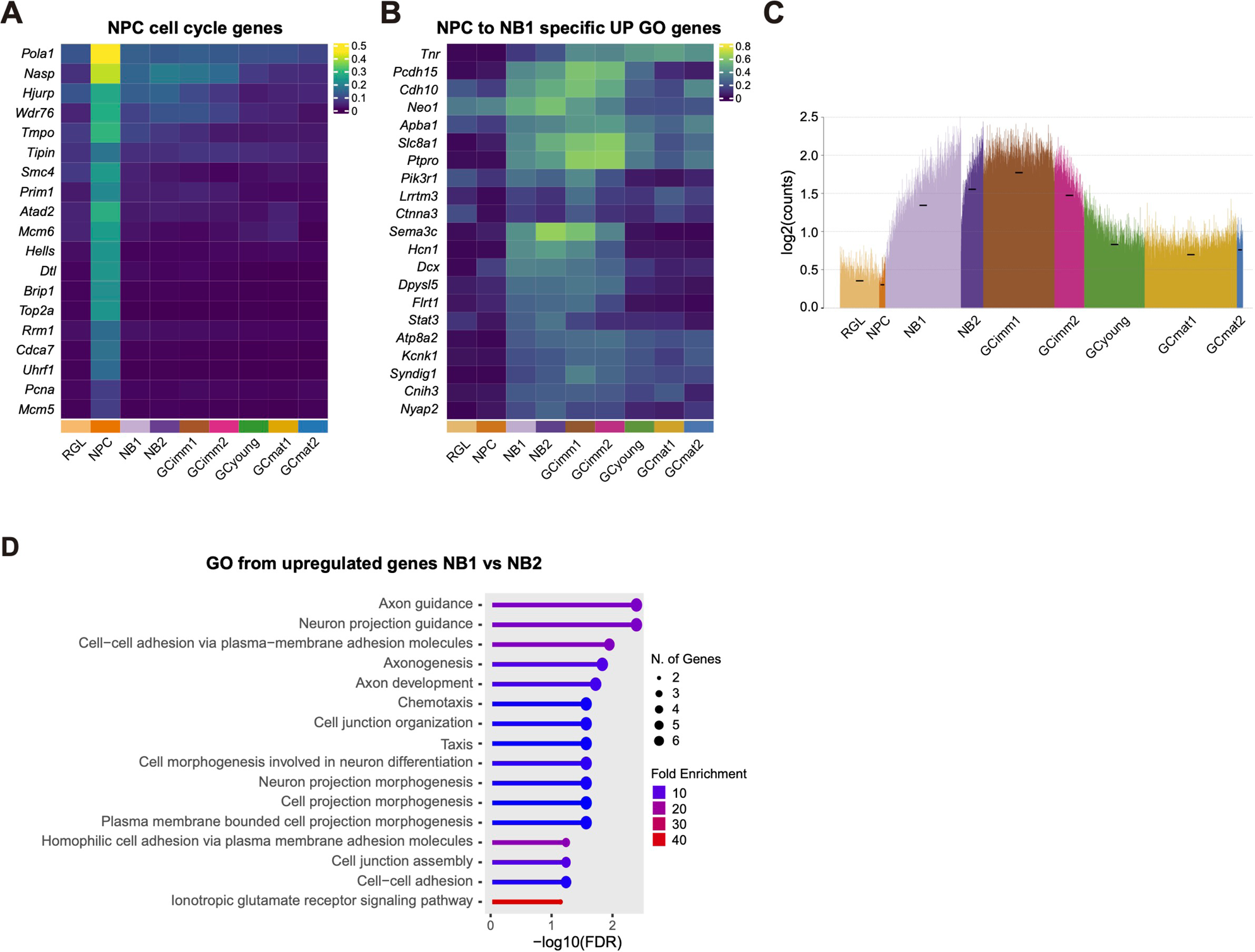
Molecular signatures of early stages of adult neurogenesis. **(A)** Heatmap showing the mean of row-wise normalized expression of canonical cell-cycle markers across clusters. Pseudocolor scale on the right denotes mean expression level. **(B)** Heatmap showing the mean of row-wise normalized expression of top 20 specific upregulated DEGs across clusters. These genes are downregulated after GCimm2. **(C)** Spike plot displaying the mean expression DEGs shown in (B). Nuclei were arranged according to their pseudotime. Black dashes show the mean expression for each cluster. **(D)** Top GO biological processes for the enrichment analysis of DEGs in the transition from NB1 to NB2. FDR cutoff = 0.05 (using ShinyGO 0.77). All data in the figure correspond to dataset 1.

**Figure S6.**
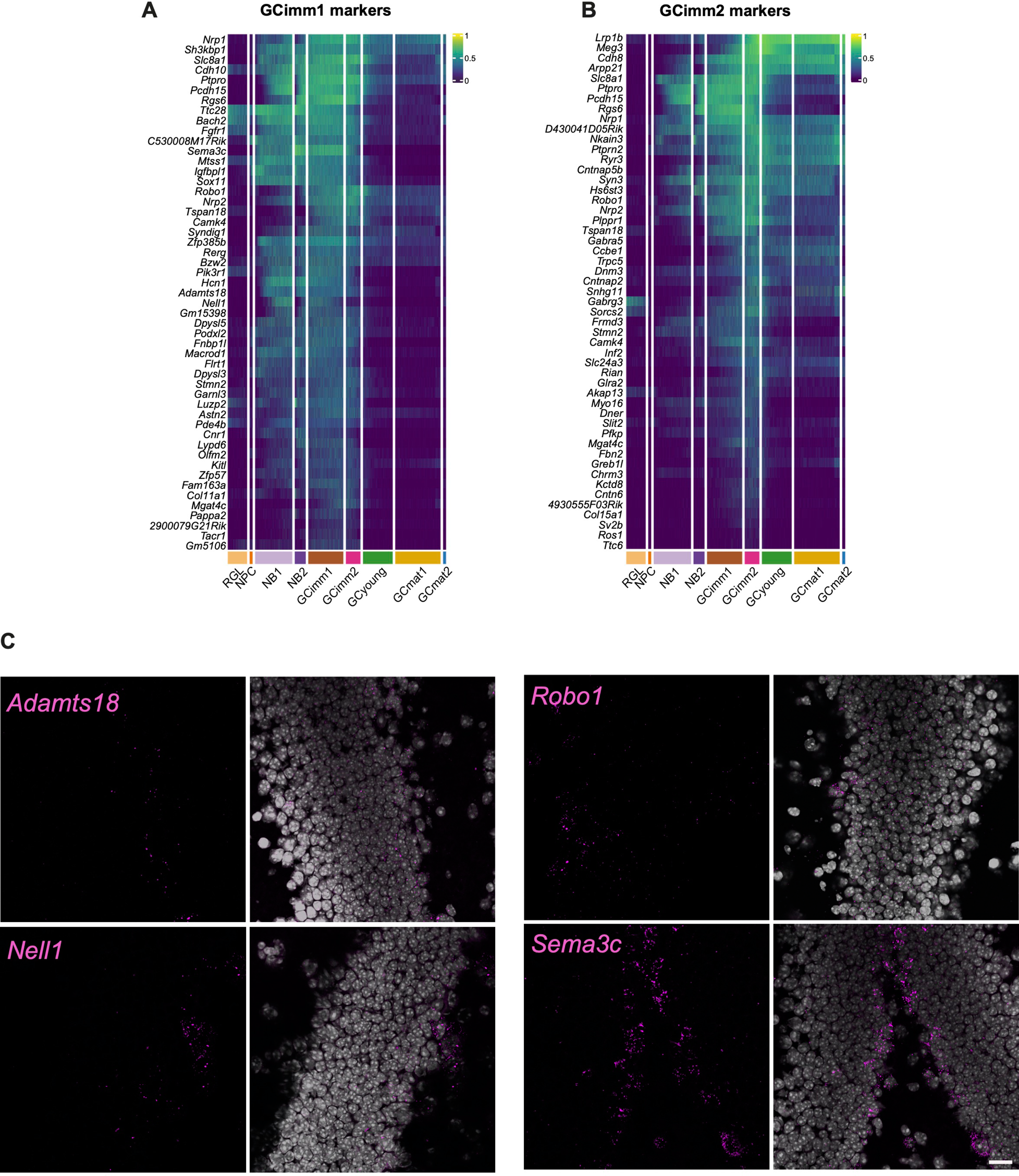
Spatial validation of GCimm1 and GCimm2 markers by RNA in situ hybridization. **(A,B)** Heatmaps showing the row-wise normalized expression of marker genes for GCimm1 and GCimm2 across clusters (top 50 genes). Marker genes were selected as described in the Methods section. Pseudocolor scales on the right denote mean expression level. Data correspond to dataset 1. **(C)** *In situ* hybridization of dentate gyrus sections from 6/7-week-old mice. Example single plane confocal images. Immature-stage markers were localized near the subgranular zone. RNA probes are shown in magenta. DAPI (grey) was used to label cell nuclei. Scale bar, 20 μm.

**Figure S7.**
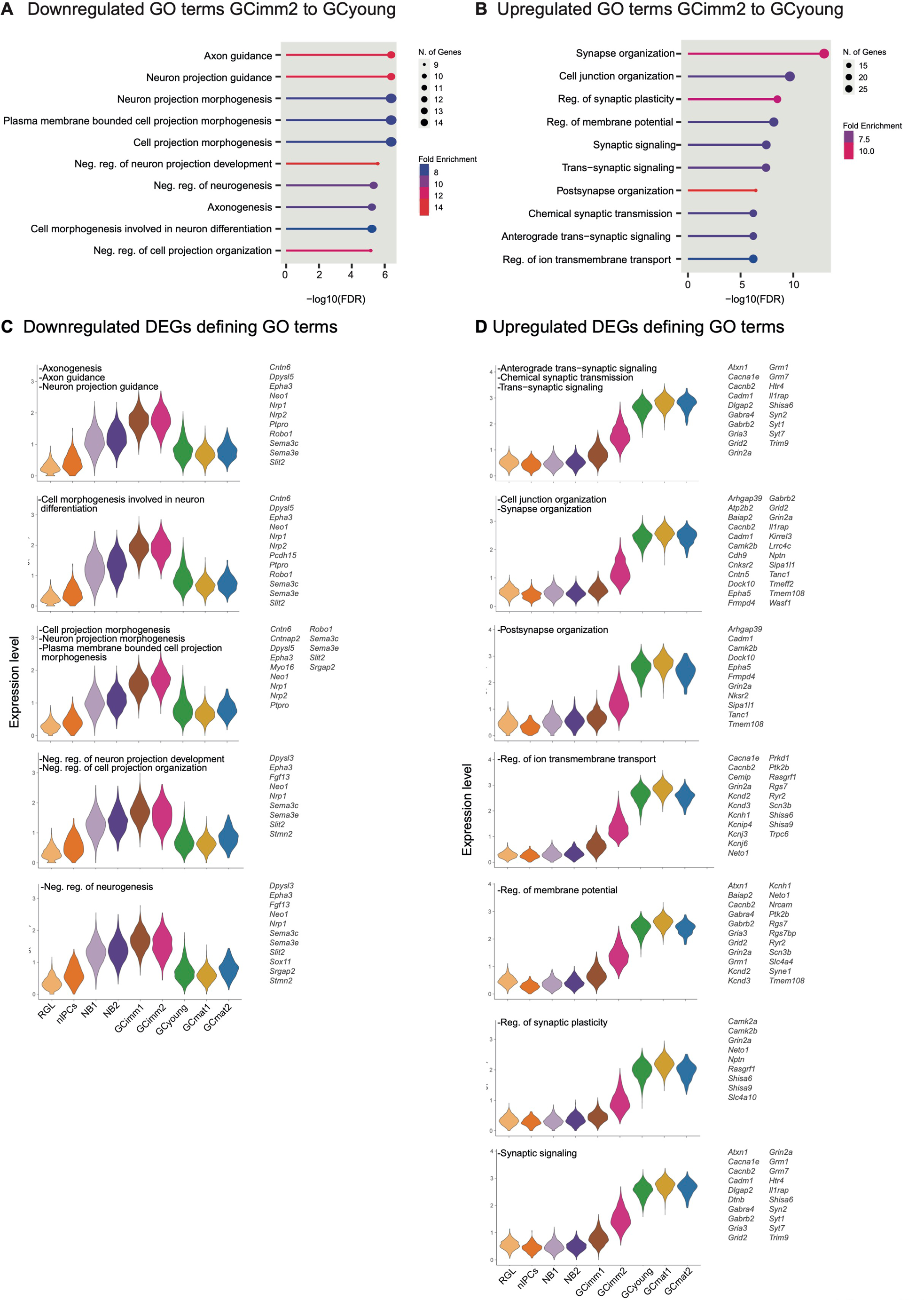
Switch in biological processes during the exit from the immature state. **(A, B)** GO biological processes are shown for the enrichment analysis of the top-50 DEGs in the transition from GCimm2 to GCyoung. FDR cutoff = 0.05 (using ShinyGO 0.77). **(C, D)** Violin plots showing the mean expression levels of DEGs for downregulated **(C)** and upregulated GO terms **(D)** for the defined clusters. All data in the figure correspond to dataset 1. DEGs are listed in **Table S2.**

**Figure S8.**
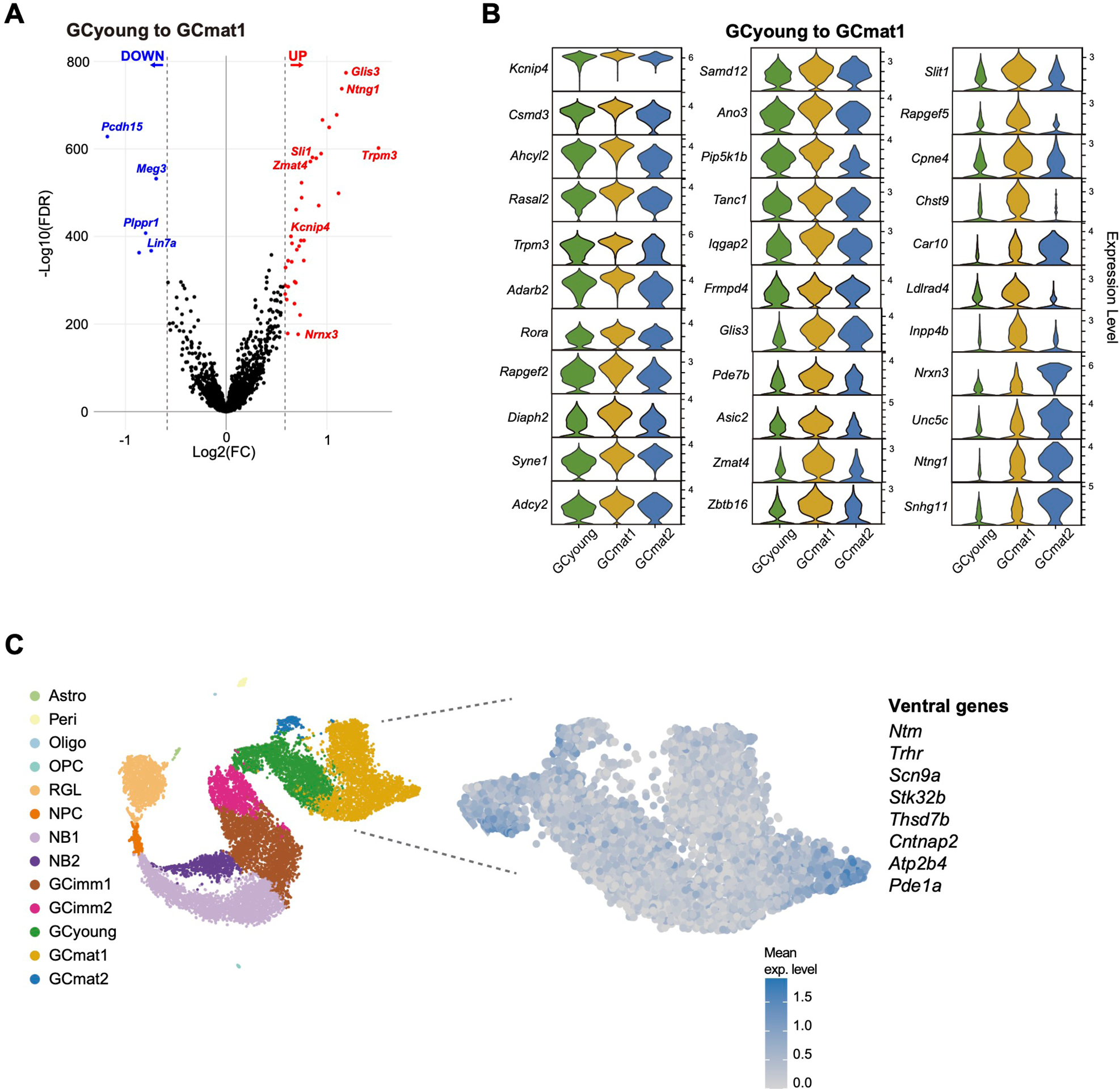
GCyoung and GCmat1 displays subtle transcriptomic differences and share nuclei with ventral signature. **(A)** Volcano plot showing differential expression analysis (FC ≥ 1.5 or ≤ −1.5 and FDR ≤ 0.05) displaying relevant gene examples. All DEGs are listed in **Table S2. (B)** Violin plots showing the expression level of upregulated DEGs of the GCyoung to GCmat1 transition. **(C)** t-SNE visualization for identified clusters (left panel). Enlarged plot section displaying the GCyoung, GCmat1 and GCmat2 clusters, depicting the mean expression of genes correlated with ventral localization: *Ntm, Trhr, Scn9a, Stk32b, Thsd7b, Cntnap2, Atp2b4 and* Pde1a. All data in the figure correspond to dataset 1.

**Figure S9.**
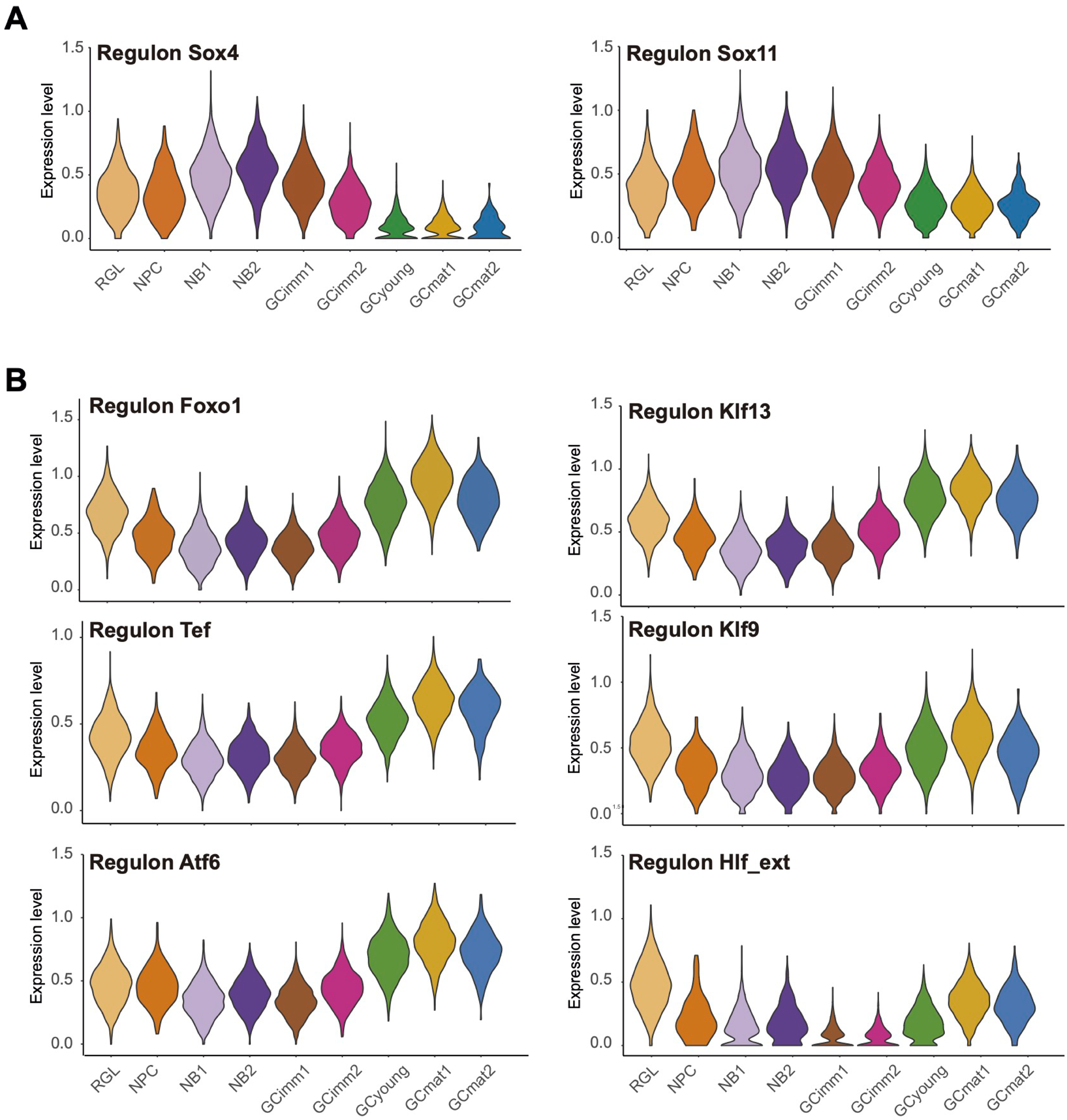
Trajectory of regulons along the different clusters. Violin plots displaying the mean expression of transcripts that compose the indicated regulons (dataset 1).

## ADDITIONAL SUPPLEMENTAL MATERIAL (Excel files)

**Table S1. DATASET 1.** Top 100 genes with highest variability, corresponding to Figure 1E.

**Table S2. DATASET 1.** Differentially expressed genes for cluster transitions in dataset 1, corresponding to Figs. 3A; 4B,C,E; 5B-D; S8A.

**Table S3. DATASET 2.** Differentially expressed genes for cluster transitions in dataset 2, corresponding to Figs. S4A.

**Table S4. DATASET 1.** Ion channels, neurotransmitter receptors, and molecules involved in synaptic transmission, corresponding to Figure 5E.

**Table S5. DATASET 2.** Regulons with differential expression between immature and mature aCGs, corresponding to Figure 7A.

